# Adaptor protein Bbc1 regulates localization of Wsp1 and Vrp1 during endocytic actin patch assembly

**DOI:** 10.1101/389015

**Authors:** Cameron MacQuarrie, MariaSanta Mangione, Robert Carroll, Michael James, Kathleen L. Gould, Vladimir Sirotkin

**Author notes:** Corresponding author: V. Sirotkin.

## Abstract

Arp2/3 complex-nucleated branched actin networks provide the force necessary for endocytosis. The Arp2/3 complex is activated by Nucleation Promoting Factors (NPFs) including the *Schizosaccharomyces pombe* proteins WASp Wsp1 and myosin-1 Myo1. There are >40 known yeast endocytic proteins with distinct spatial and temporal localizations and functions; however, it is still unclear how these proteins work together to drive endocytosis. We used quantitative live cell imaging to determine the function of the uncharacterized *S. pombe* protein Bbc1. We discovered Myo1 interacts with and recruits Bbc1 to sites of endocytosis. Bbc1 competes with verprolin Vrp1 for Myo1 binding, thus releasing Vrp1 and its binding partner Wsp1 from Myo1. Normally Myo1 remains at the base of the endocytic invagination and Vrp1-Wsp1 internalize with the endocytic vesicle; however, in the absence of Bbc1, a portion of Vrp1-Wsp1 remains with Myo1 at the base of the invagination and endocytic invaginations are twice as long. We propose that Bbc1 disrupts a transient Myo1-Vrp1-Wsp1 interaction and limits Arp2/3 complex-nucleation of actin branches at the plasma membrane.

---

**Abbreviations**
WASp: Wiskott-Aldrich Syndrome protein
NPF: Nucleation Promoting Factor
CR: contractile ring
SPB: Spindle Pole Body
SPF: seconds per frame

## INTRODUCTION

Clathrin Mediated Endocytosis (CME) is an essential process conserved across eukaryotes, which allows for the internalization of external materials into a cell. This process is highly regulated with >40 proteins known to assemble into patch structures at sites of endocytosis in a consistent spatial and temporal pattern (Goode et al., 2015; Kovar et al., 2011; McMahon and Boucrot, 2011; Weinberg and Drubin, 2012). How these proteins coordinate their activities to ensure reproducible endocytic patch assembly and efficient endocytosis is an important open question in the field. Although many of the endocytic patch proteins are conserved among species, some are specific to animals or yeast. Bbc1/Mti1 is a yeast specific protein, which localizes to sites of endocytosis and is thought to function in regulating actin assembly (Kaksonen et al., 2005; Mochida et al., 2002; Picco et al., 2018; Rodal et al., 2003; Sun et al., 2006).

To help overcome membrane tension in yeast and animal cells, endocytosis relies upon the assembly of actin into structures called actin patches, which provide the scaffold and force needed for efficient membrane invagination and scission (Aghamohammadzadeh and Ayscough, 2009; Basu et al., 2014; Boulant et al., 2011; Engqvist-Goldstein and Drubin, 2003; Goode et al., 2015; Kaksonen and Roux, 2018; Kaksonen et al., 2006). The branched actin network at endocytic sites (Collins et al., 2011; Rodal et al., 2005; Young et al., 2004) is assembled by the Arp2/3 complex upon its activation by Nucleation Promoting Factors (NPFs) (Higgs and Pollard, 2001). Multiple NPFs are present at endocytic sites in fission yeast *Schizosaccharomyces pombe* (Basu and Chang, 2011; Carnahan and Gould, 2003; Kovar et al., 2011; Sirotkin et al., 2005; Sirotkin et al., 2010) including Wiskott-Aldrich Syndrome protein (WASp) homolog Wsp1, myosin-1 Myo1, EPS15 homolog Pan1, and SPIN90 homolog Dip1. Homologs of these proteins are also present in budding yeast *Saccharomyces cerevisiae*, including WASp Las17 and myosin-1 Myo3/5 (Boettner et al., 2012; Goode et al., 2015; Weinberg and Drubin, 2012). NPFs share the ability to stimulate Arp2/3 complex but differ in the strength and mechanism of Arp2/3 complex activation and have additional distinct functions (Galletta et al., 2008; Sirotkin et al., 2005; Sun et al., 2006; Wagner et al., 2013). Specifically, in both budding and fission yeast, WASp is the strongest Arp2/3 complex activator and is considered the primary NPF driving actin assembly for membrane deformation, while myosin-1 is a weak activator that is thought to contribute force primarily via its actin-dependent motor activity (Basu et al., 2014; Berro et al., 2010; Sirotkin et al., 2005; Sun et al., 2006). These diverse NPFs collaborate during endocytosis. In budding yeast, Las17 and Myo3/5 form a WASP/myosin-1 module (Kaksonen et al., 2005; Weinberg and Drubin, 2012) that also includes a homolog of WASp Interacting protein (WIP) verprolin Vrp1 and F-BAR protein Bzz1, both of which regulate NPF activity in budding and fission yeast (Arasada and Pollard, 2011; Sirotkin et al., 2005; Soulard et al., 2002; Sun et al., 2006). A similar collaboration exists in fission yeast where Myo1 and Wsp1 interact directly (Carnahan and Gould, 2003) or indirectly via Vrp1 that is recruited to patches by Wsp1 but can also bind Myo1 and stimulate its NPF activity (Sirotkin et al., 2005). Myo1 localizes to patches independently of Wsp1, but accumulation of Wsp1 and Vrp1 is reduced in the absence of Myo1 (Sirotkin et al., 2005) (Our unpublished observations).

The localization and activity of NPFs is regulated to ensure proper actin polymerization. One proposed regulatory component of the budding yeast WASP/myosin-1 module of endocytosis is Bbc1/Mti1 (Kaksonen et al., 2005; Weinberg and Drubin, 2012). Bbc1 was originally identified in *S. cerevisiae* as a binding partner of myosin-1 Myo5 (Mochida et al., 2002) and was shown to mildly inhibit Myo5 NPF activity *in vitro* (Sun et al., 2006). However, Bbc1 also interacts with Las17 (Tong et al., 2002). Consistent with this interaction, Bbc1, along with Myo5, is recruited to Las17-coated beads but not to Myo5 in comet tails in *in vitro* re-constitution assays (Michelot et al., 2010). Moreover, a pioneering study by Rodal et al. (2003) demonstrated that Bbc1, in cooperation with adaptor protein Sla1, inhibits Las17 NPF activity *in vitro*. This work led to a widely accepted model for Bbc1 and Sla1 as dual Las17 inhibitors (Weinberg and Drubin, 2012). Supporting an inhibitory function for Bbc1, *bbc1*Δ *sla1*Δ and, to a lesser extent, *bbc1*Δ cells have enhanced actin patch assembly (Kaksonen et al., 2005; Picco et al., 2018) The *bbcl*Δ cells are also characterized by deeper internalizations of endocytic structures (Kaksonen et al., 2005; Picco et al., 2018); however, this depends on the NPF activity of Myo5 but not Las17 (Sun et al., 2006), consistent with Bbc1 identification as a Myo5 ligand. The behavior and precise localization of Bbc1 at endocytic sites further suggest that Bbc1 interaction with Myo5 may be important for additional, yet to be discovered functions of Bbc1 in cells, beyond direct inhibition of Las17 NPF activity. Specifically, *S. cerevisiae* Bbc1 is recruited to actin patches with timing similar to Myo3/5 (Kaksonen et al., 2005) and while immuno-EM localized Bbc1 near both Las17 and Myo3/5 *in vivo* (Idrissi et al., 2008), a recent super-resolution microscopy study placed Bbc1 closer to Myo3/5 than Las17 (Mund et al., Nov. 15, 2017). Yet, the significance of the Bbc1 interaction with myosin-1, other than the ability of *S. cerevisiae* Bbc1 to inhibit Myo5 NPF activity *in vitro* (Sun et al., 2006), remains largely unknown.

Interestingly, the localization pattern of key NPFs differs in budding and fission yeasts. In budding yeast, Myo3/5 and Las17 remain at the base of the invagination and do not internalize with the endocytic vesicle (Kaksonen et al., 2005). In contrast, *S. pombe* Myo1 and Wsp1 show distinct localization and behavior: Myo1 does not internalize with the actin patch and remains at the base of the endocytic invagination while Wsp1 internalizes with the patch and separates from Myo1 (Sirotkin et al., 2005). This spatial separation of Myo1 and Wsp1 in *S. pombe* makes it possible to determine whether regulators of actin patch assembly function via association with Myo1 or Wsp1 by following protein dynamics in live cells.

We detected a protein homologous to budding yeast Bbc1/Mti1 in pull-downs with contractile ring (CR) proteins Cyk3 (Roberts-Galbraith et al., 2010) and Fic1, piquing our interest in this thus far uncharacterized *S. pombe* protein. Here, we take advantage of distinct behaviors of Myo1 and Wsp1 in *S. pombe* to reveal a novel mechanism for how Bbc1 regulates actin assembly based on a competition between Bbc1 and Vrp1-Wsp1 for binding the Myo1 tail.

## RESULTS

### Bbc1 Localizes to the Cell Division Site but not the Contractile Ring

The protein SPAC23A1.17/Mti1/Bbc1 was identified by mass spectrometry (MS) in affinity purifications of the CR proteins Cyk3 (Roberts-Galbraith et al., 2010) and Fic1 (Table S1), prompting us to investigate the localization of this uncharacterized *S. pombe* protein in dividing cells (Figure 1). Bbc1-mNeonGreen (mNG) was absent from the CR labeled with mCherry-Cdc15 but was present at the division site in two types of structures: dynamic cortical puncta, which were also present at cell tips in interphase and at the onset of cell division (Figure 1A), and stable puncta, which emerged upon completion of CR constriction (Figure 1B). Imaging cells expressing both Bbc1-mNG, spindle pole body (SPB) marker Sid4-mNG, and CR marker mCherry-Cdc15 revealed that Bbc1-mNG arrives to the division site after CR formation, 20.9 ± 2.7 minutes after the onset of mitosis defined by the separation of SPBs (Figures 1, B and C). This is similar to the timing of appearance of endocytic actin patches at the division site (Wu et al., 2006). This timing was further confirmed by end-on imaging (Figure 1D) of *S. pombe* expressing Bbc1-mNG and the CR component Rlc1-mCherry (Laporte et al., 2011; Le Goff et al., 2000; Wu et al., 2006), which also revealed that Bbc1, like other actin patch components (Wu et al., 2006), arrives late to the division site and does not constrict with the CR. Imaging cells expressing Bbc1-mGFP and endocytic actin patch marker Fim1-mCherry revealed that the dynamic cortical puncta of Bbc1 at the division site are indeed actin patches and the stable puncta are not (Figure 1, E and F).

**Figure 1:**
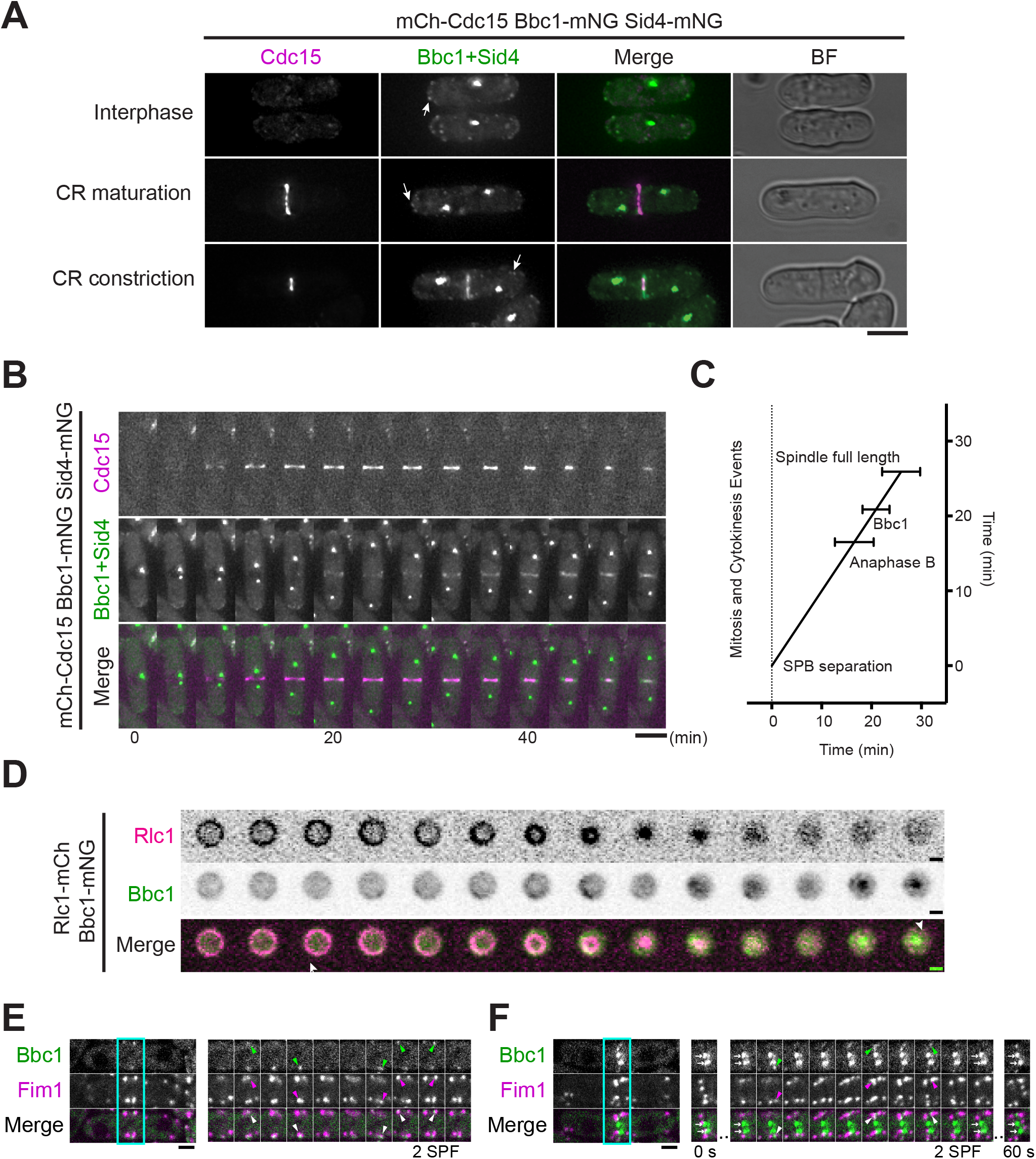
Bbc1 localizes to the cell division site but not the CR. (A) Representative live-cell images of mCherry-Cdc15 (magenta), Bbc1-mNeonGreen (mNG) and Sid4-mNG (green) at different cell cycle stages. Arrows indicate Bbc1 cortical puncta. Scale bar, 5 μm. (B) Images at 4-minute intervals from a representative movie of cells expressing mCherry-Cdc15 (magenta), Bbc1-mNG and Sid4-mNG (green). Scale bar, 5 μm. (C) Time line showing detection of Bbc1 at the division site. Time 0 was defined as the time of SPB separation. The mean time of each event is plotted. Error bars represent S.E.M. The time of the onset of anaphase B was determined to be 16.5 +/− 3.9 minutes and the time of the maximum SPB separation was determined to be 25.9 +/− 3.9 minutes. Bbc1 arrived at the division site 20.9 +/− 2.7 minutes after SPB separation. N=23 cells. (D) End-on images at 4-minute intervals of cells expressing the CR marker Rlc1-mCherry (magenta) and Bbc1-mNG (green). Scale bars, 2 μm. (E, F) Time lapse images of Bbc1-mGFP (green) and Fim1-mCherry (magenta) at the division site (boxed in teal) at the (E) beginning and (F) end of CR constriction. Images were acquired at the rate of 2 seconds per frame (SPF) for 60 seconds. Arrowheads mark dynamic patches and arrows mark stable puncta. Scale bars, 2 μm.

To gain insight into the physiological role of Bbc1, we deleted *bbc1^+^* and examined genetic interactions of *bbc1*Δ and sensitivity of *bbc1*Δ cells to a variety of stresses. The *bbc1*Δ cells were sensitive to benomyl and latrunculin A (Supplemental Figure S1A) and showed negative interactions with *fim1*Δ, *myo1*Δ, *zds1*Δ, and *mid1*Δ (Supplemental Figure S1B), implicating Bbc1 in endocytosis, cell wall integrity, and cytokinesis (Chang et al., 1996; Sirotkin et al., 2005; Skau and Kovar, 2010; Sohrmann et al., 1996; Yakura et al., 2006). However, *bbc1*Δ cells were morphologically normal and had no apparent cytokinesis defects, and Bbc1-overexpressing cells had a normal septation index (Supplemental Figure S1C). Given that *S. cerevisiae* Bbc1 has been identified as a myosin-1 ligand (Mochida et al., 2002), the fact that *S. pombe* Bbc1 pulls down actin patch components Myo1, Pan1, and Cdc15 in affinity purifications (Table S1), and that Bbc1 localizes to actin patches in both *S. cerevisiae* (Mochida et al., 2002) and *S. pombe* (this study), we then investigated the role of *S. pombe* Bbc1 in endocytic actin patches.

### Bbc1 Localizes to the Base of the Endocytic Invagination in a Myo1-dependent manner

We confirmed by co-localization with actin patch components Fim1 and Pan1 that in interphase cells, as during cell division, Bbc1 localizes to both actin patches and actin-free puncta (Figure 2, A and B, and Supplemental Figure S2A). During interphase, Bbc1 in actin patches also co-localizes with Cdc15 (Figure 2C), which localizes to actin patches and stable puncta in interphase (Supplemental Figure S2B) but leaves actin patches for CR during cell division (Arasada and Pollard, 2011; Carnahan and Gould, 2003; Wu et al., 2006).

**Figure 2:**
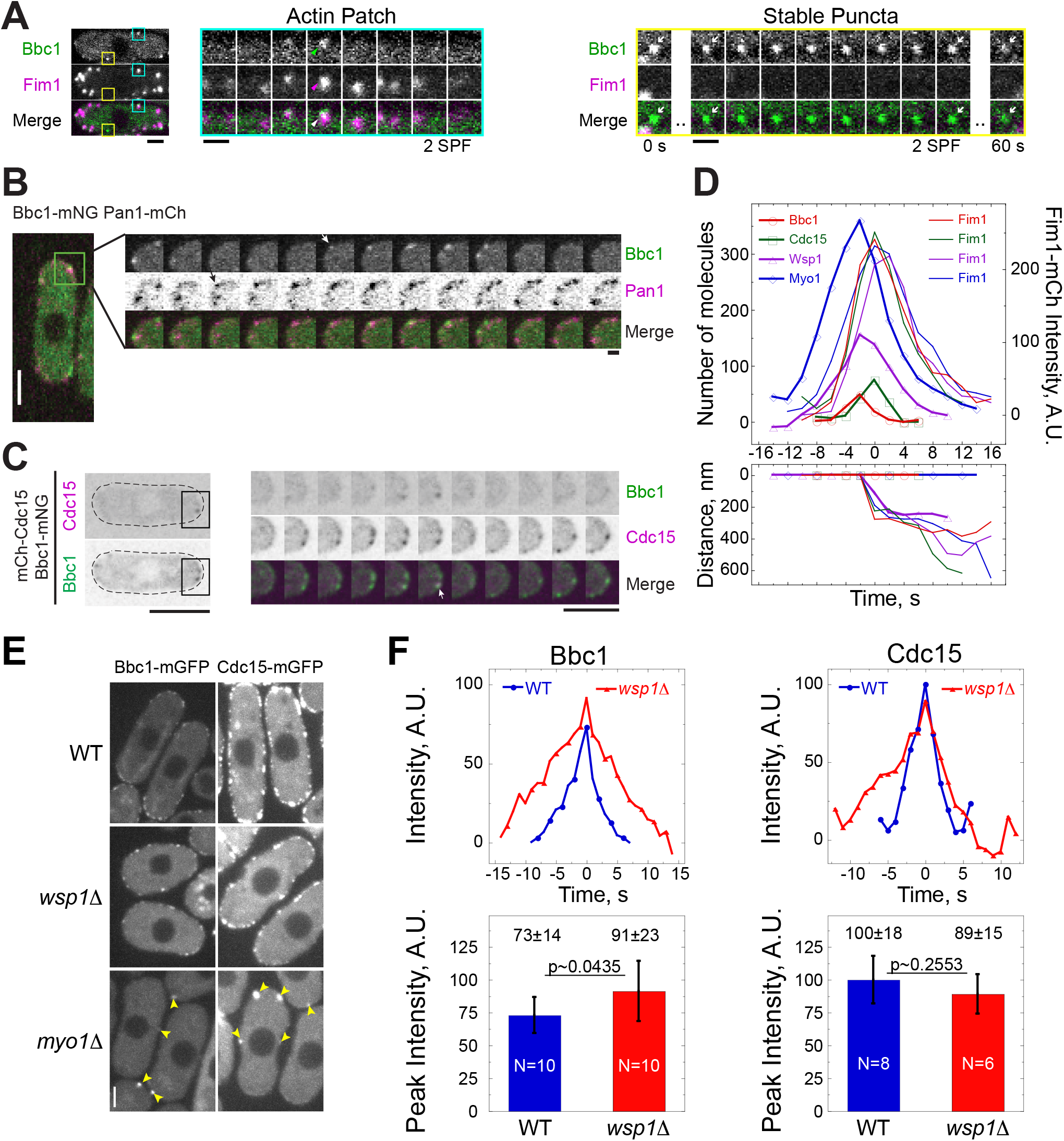
Bbc1 localizes to the base of the endocytic invagination in a Myo1-dependent manner. (A-C) Time lapse images of (A) Bbc1-mGFP (green) and Fim1-mCherry (magenta), (B) Bbc1-mNG (green) and Pan1-mCherry (magenta), and (C) Bbc1-mNG (green) and Cdc15-mCherry (magenta) at 2-second (A, B) or 1-second (C) intervals in boxed regions of single live interphase cells (left panels). In (A), arrowheads indicate an actin patch in a region boxed in teal and arrows indicate a stable punctum in a region boxed in yellow. In (B) and (C), arrows mark protein appearances in endocytic actin patches. Scale bars, (A) cell panel, 2 μm, montages, 1 μm; (B) cell panel, 5 μm, montages, 2 μm; (C) cell panel and montages, 5 μm. (D) Average time courses of the number of molecules of mGFP-tagged Bbc1, Cdc15, Wsp1, and Myo1, Fim1-mCherry fluorescence intensity, and distance traveled by each marker in endocytic actin patches in cells expressing Bbc1-mGFP (red circles) and Fim1-mCherry (red, no symbol), Cdc15-mGFP (green squares) and Fim1-mCherry (green, no symbol), mGFP-Wsp1 (purple triangles) and Fim1-mCherry (purple, no symbol), or mGFP-Myo1 (blue diamonds) and Fim1-mCherry (blue, no symbol). Images in a medial plane were acquired at 2-second intervals. Intensity and position of each marker in 6-9 actin patches were tracked. Individual time courses were aligned to the start of Fim1-mCherry patch movement (t=0), averaged at each time point, and mGFP intensities were converted to the number of molecules (see Materials and Methods) based on previous measurements (Sirotkin et al., 2010). (E) Maximum intensity projections of time series of images in a single confocal section through the middle of the WT, *wsp1Δ*, and *myo1Δ* cells expressing Bbc1-mGFP (left) or Cdc15-mGFP (right). Images were acquired at 1-second intervals for 60 seconds. Scale bar, 2 μm. Yellow arrowheads indicate stable puncta. Also see Figure S2. (F) Average time courses of intensities and bar graphs of peak intensities of Bbc1-mGFP and Cdc15-mGPF in endocytic patches in WT (blue) and *wsp1Δ* (red) cells. Time courses for individual patches were aligned to peak patch intensity (t=0) and averaged at each time point. P values represent statistical significance as determined by t-test. N indicates number of patches analyzed. Error bars represent SD.

Quantitative imaging of strains combining Fim1-mCherry with mGFP-tagged Bbc1 or other patch proteins provided detailed accounting of Bbc1 dynamics in patches relative to Myo1, Wsp1, and Cdc15, patch markers with known dynamics (Arasada and Pollard, 2011; Sirotkin et al., 2010). We determined the time courses of the numbers of molecules of these proteins in actin patches (Figure 2D and Supplemental Figure S2, C-G) based on previously measured numbers of actin patch proteins (Sirotkin et al., 2010) (see Materials and Methods). Time courses of the numbers of molecules were aligned to the time when patches marked by Fim1-mCherry first moved (time zero), which corresponds to the initiation of actin patch internalization. *S. pombe* Bbc1 arrived to actin patches 4-6 seconds before the start of actin patch internalization, reached peak number of 48 molecules at −2 seconds, and quickly dissipated 2-6 seconds after time zero. Cdc15 showed similar behavior, appearing at −4 to −2 seconds, peaking at 74 molecules at time zero, and disappearing 2-4 seconds later, comparable to previous measurements (Arasada and Pollard, 2011). Both Myo1 and Wsp1 are present in excess over Bbc1 and Cdc15, reaching peaks of 359 and 155 molecules, respectively at −2 seconds. In contrast to Wsp1, which in *S. pombe* internalizes with the actin patch, neither Bbc1 nor, as previously reported (Arasada and Pollard, 2011), Cdc15 internalized with actin patches. This behavior of Bbc1 and Cdc15 resembles that of Myo1, suggesting Bbc1 and Cdc15 possibly function with Myo1 at the base of the endocytic invagination.

Since Bbc1 in *S. cerevisiae* was initially identified as a myosin-1 Myo5 ligand (Mochida et al., 2002) but later shown to also bind and inhibit *S. cerevisiae* WASp Las17 (Rodal et al., 2003), we tested whether *S. pombe* Myo1 and Wsp1 are important for Bbc1 localization to patches in *S. pombe*. We examined the genetic dependencies of Bbc1 localization to dynamic cortical patches using Cdc15, which is known to depend on Myo1 for localization to actin patches (Arasada and Pollard, 2011), as a control (Figure 2, E and F). We found that both Bbc1-mGFP and Cdc15-mGFP failed to localize to dynamic cortical patches in *myo1*Δ cells (Figure 2E). Instead, in interphase *myo1*Δ cells, Bbc1-mGFP and Cdc15-mGFP were observed exclusively in stable cortical puncta (Supplemental Figure S2, A and B). In contrast, in the absence of Wsp1, both Bbc1-mGFP and Cdc15-mGFP continued to localize in dynamic patches (Figure 2E), albeit with slower dynamics (Figure 2F) typical for endocytic patches in *wsp1*Δ cells (Basu et al., 2014). Interestingly, while Cdc15 accumulated in patches to the same level in *wsp1*Δ as in wild-type cells, recruitment of Bbc1 to patches in *wsp1*Δ cells was enhanced by ~25% (Figure 2F). Thus, Bbc1 is recruited to patches by Myo1 and may compete with other ligands for binding Myo1.

### Myo1 Recruits Bbc1 through the TH2 and SH3 domains

To determine which part of Myo1 is responsible for recruiting Bbc1 and Cdc15 to patches, we examined the ability of Bbc1 and Cdc15 to localize to patches marked by Fim1-mCherry in strains featuring a series of Myo1 tail truncations using two-color imaging (Figure 3A and Supplemental Figure S3). We uncovered that the Myo1 TH2 and SH3 domains are both required for Bbc1 patch recruitment (Figure 3A). In contrast, the Myo1 TH2 and SH3 domains contribute but are not required for Cdc15 recruitment as Cdc15 continued to localize to some of the patches, albeit at decreased levels, when the Myo1 TH2 domain, SH3 domain, or both were deleted (Supplemental Figure S3). This is consistent with previous data, which indicated that the Myo1 TH1 and TH2 domains contribute to an interaction with Cdc15 (Carnahan and Gould, 2003), but in contrast to a previous report of an apparent loss of Cdc15 from patches in *myo1*Δ*23A* mutant lacking the Myo1 TH2, SH3, and LCA domains (Arasada and Pollard, 2011).

**Figure 3:**
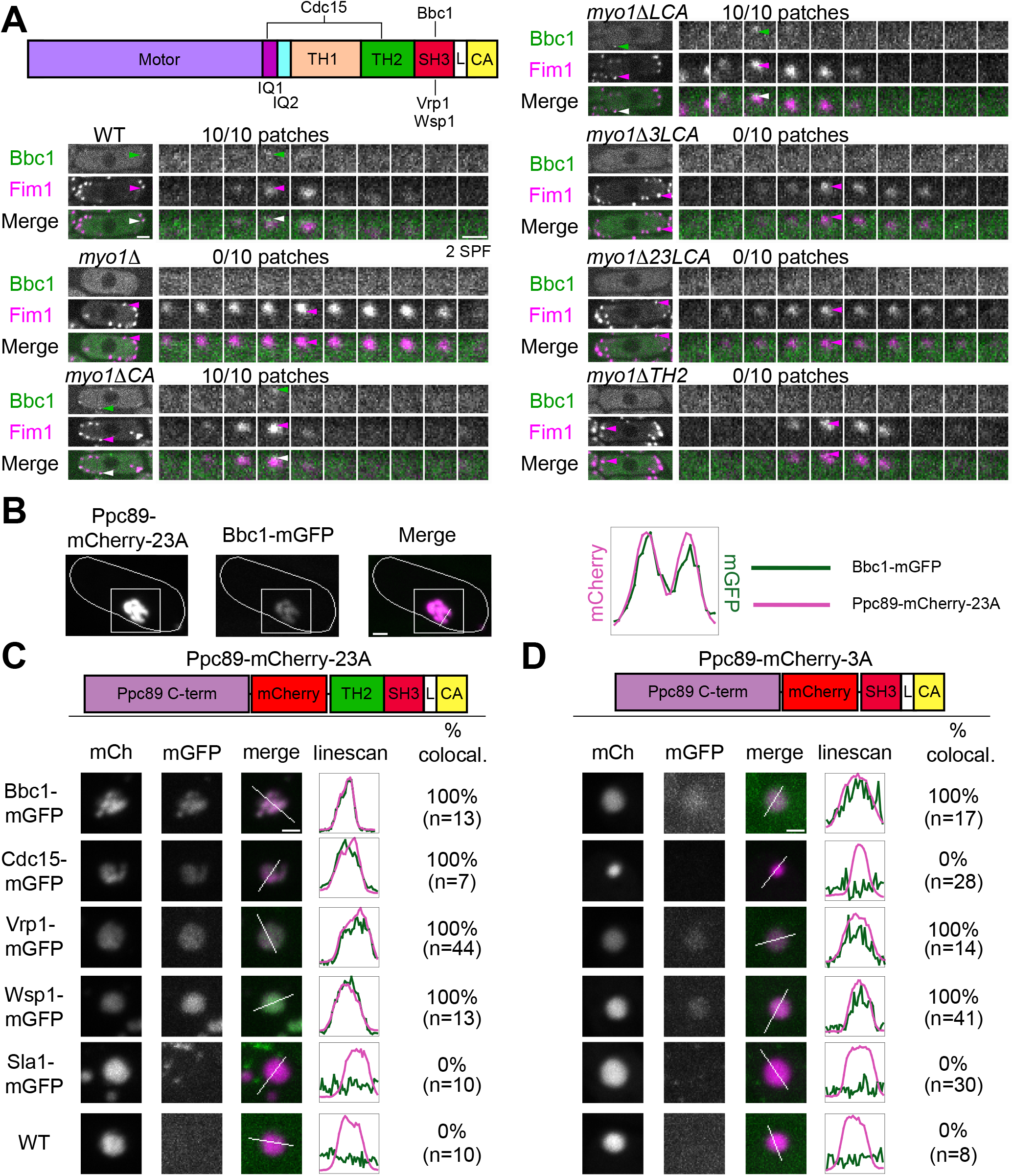
The Myo1 SH3 domain is necessary to recruit Bbc1. (A) A schematic diagram of the Myo1 protein domains with relevant binding partners and example montages of co-localization analysis of Bbc1-mGFP (green) and Fim1-mCherry (magenta) in patches in WT and indicated Myo1 tail domain deletion strains. Montages show the protein localization dynamics at intervals of 2 SPF in patches indicated by arrowheads on montages and whole cell images on the left. Numbers show the fraction of Fim1 patches that also contained Bbc1. Scale bars, whole cells, 2 μm; montages, 1 μm. (B) Images of a fission yeast cell (outlined) showing co-localization of a Ppc89-mCherry-23A (magenta) aggregate and Bbc1-mGFP (green). A line scan through the aggregate confirms co-localization. (C, D) Representative images and line scans from co-localization analysis of depicted (C) Ppc89-mCherry-23A or (D) Ppc89-mCherry-3A protein constructs expressed off of plasmid in WT cells or in cells expressing mGFP-tagged Bbc1, Cdc15, Vrp1, Wsp1 and Sla1 from endogenous loci. The graphs show single line scans of mCherry and mGFP intensities along the lines across aggregates indicated by white lines on merged mCherry (magenta) and mGFP (green) images. Numbers indicate percent co-localization; n indicates number of aggregates analyzed for each condition. Scale bars, 1 μm.

To confirm these localization dependencies, we fused the C-terminal domain of the SPB protein Ppc89 (Rosenberg et al., 2006) to mCherry and Myo1 tail fragments consisting of the TH2, SH3, and LCA domains (Ppc89-mCherry-23A) or the SH3 and LCA domains (Ppc89-mCherry-3A). These constructs were introduced into cells expressing mGFP-tagged Bbc1, other known Myo1 tail ligands Vrp1, Wsp1, and Cdc15 (Carnahan and Gould, 2003; Sirotkin et al., 2005), or Sla1, which is not known to interact with Myo1. Rather than localizing to SPBs as might have been expected, both Ppc89-mCherry-23A and −3A constructs formed aggregates. These were still useful, however, to determine whether mGFP-tagged proteins were recruited by mCherry-marked Myo1 fragments (Figure 3B). With the exception of Sla1, all examined proteins relocalized to the Ppc89-mCherry-23A puncta (Figure 3C). However, only Bbc1, Vrp1, and Wsp1 relocalized to the Ppc89-mCherry-3A puncta (Figure 3D). The binding of Cdc15 to the TH2-SH3-LCA but not SH3-LCA Myo1 construct is consistent with previously reported interaction of Cdc15 with the Myo1 TH2 domain (Carnahan and Gould, 2003) and suggests that the reduction of Cdc15 at patches in *myo1* Δ*SH3-LCA* cells could be explained by loss of indirect interactions that bridge Myo1 and Cdc15. In contrast, the Myo1 SH3 domain is sufficient for interaction with Bbc1 and together with the TH2 domain is necessary for Bbc1 localization to patches.

### Vrp1 Competes with Bbc1 for interaction with Myo1 and Helps Retain Myo1 in Patches

Since the Myo1 SH3 domain binds Vrp1 *in vitro* (Sirotkin et al., 2005) and recruits both Vrp1 and Bbc1 in our *in vivo* binding assay (Figure 3D), we tested whether Vrp1 and Bbc1 compete for access to Myo1 in cells. We measured accumulation of Bbc1-mGFP in patches in *vrp1 Δ* cells and Vrp1-mGFP in *bbc1 Δ* cells versus wild-type cells, expecting increased accumulation of Bbc1 or Vrp1 in patches in the absence of its potential competitor. Indeed, Bbc1 patch accumulation was increased by 40% in the absence of Vrp1 (Figure 4A), while Vrp1 patch accumulation was increased by 51% in the absence of Bbc1 (Figure 4B). This competition mechanism may also explain the modest 25% increase in the Bbc1 level in patches observed in *wsp1 Δ* cells (Figure 2F) since Wsp1 recruits Vrp1 to patches (Sirotkin et al., 2005).

**Figure 4:**
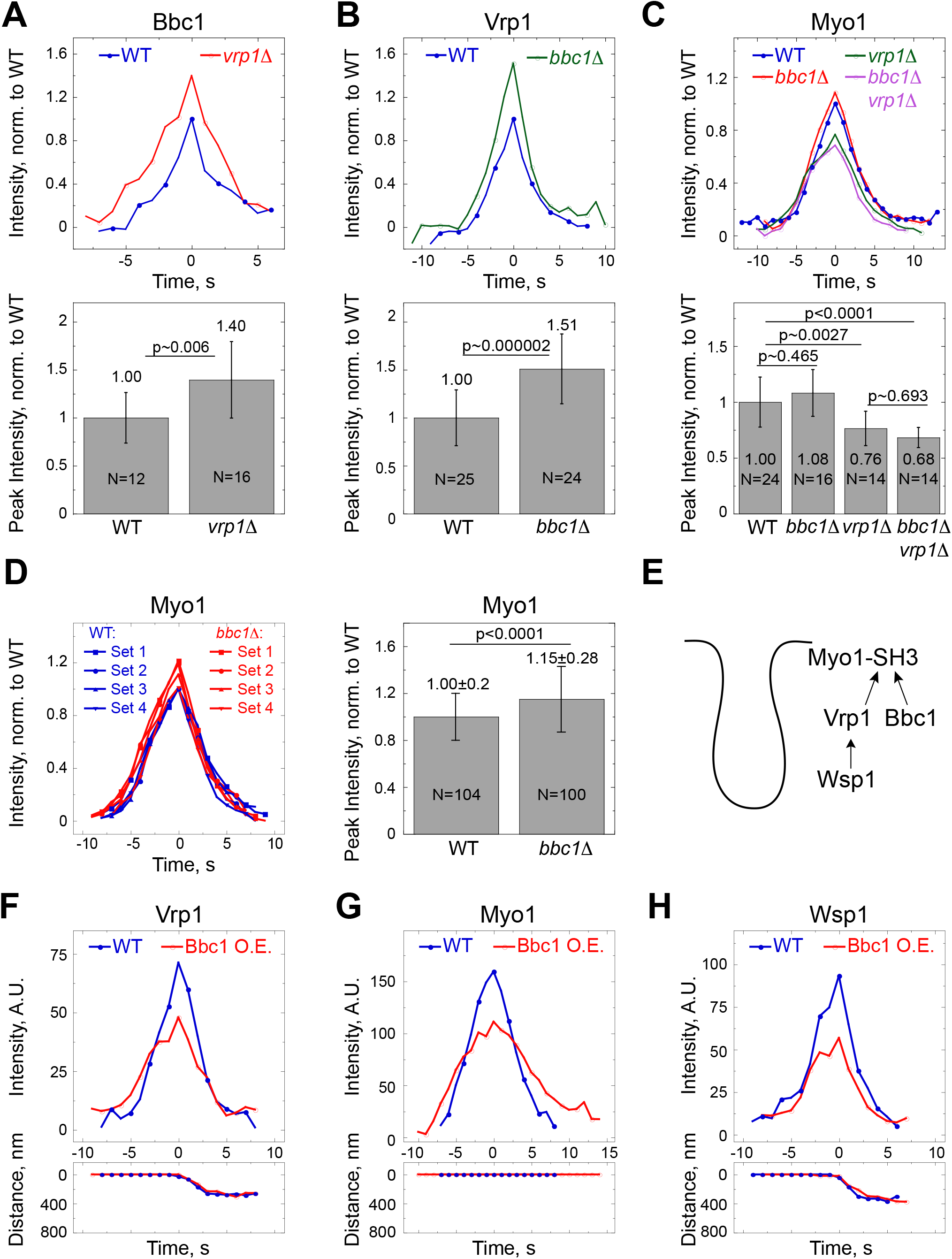
Bbc1 and Vrp1 compete for Myo1 and modulate NPF accumulation in patches. (A-D) Average time courses of normalized intensities and bar graphs of average normalized peak intensities of (A) Bbc1-mGFP in patches in WT (blue) and *vrp1Δ* (red) cells, (B) Vrp1-mGFP in patches in WT (blue) and *bbc1Δ* (green) cells, (C) mGFP-Myo1 in patches in WT (blue), *bbc1Δ* (red), *vrp1Δ* (green), *bbc1Δ vrp1Δ* (purple) cells, and (D) GFP-tagged Myo1 in patches in WT (blue) and *bbc1Δ* (red) cells. Time courses for individual patches were aligned to the time each patch reached peak intensity (t=0), averaged at each time point, and normalized to the average peak intensities in the WT cells. Myo1 patch dynamics were measured in WT (blue) or *bbc1Δ* (red) cells expressing Myo1-GFP (Sets 1 and 3), Myo1-GFP and LifeAct-mCherry (Set 2), or mGFP-Myo1 (Set 4) and normalized peak intensities for individual patches from all datasets were combined and averaged. Error bars represent SD. N indicates number of patches analyzed. P values represent statistical significance as determined by (A, B, and D) t-test or (C) ANOVA with Tukey’s Post-Hoc test. (E) A schematic showing Bbc1 competing with Vrp1 and Wsp1 for the Myo1 SH3 domain. (F-H) Average time courses of intensities and distances traveled by (F) Vrp1-mGFP, (g) mGFP-Myo1, or (H) mGFP-Wsp1 in endocytic patches in WT (blue) or Bbc1-overexpressing (O.E., red) cells. Time courses for individual patches were aligned to the time each patch reached peak intensity (t=0). N=6-8 patches per condition.

Next, we examined the effects of competition of Bbc1 or Vrp1 for Myo1 tail on Myo1 recruitment to patches by measuring Myo1 levels in patches in the absence of Bbc1, Vrp1, or both proteins (Figure 4C). Myo1 accumulation in patches was reduced by 20-30% in *vrp1*Δ and *bbc1Δ vrp1*Δ cells, suggesting that Vrp1 helps retain Myo1 in patches even though these proteins differ in their patch dynamics (Sirotkin et al., 2005). In contrast, Myo1 accumulation was mildly increased in *bbc1*Δ cells. This slight increase, 15% on average, was reproducible across multiple experiments (Figure 4D) and can be explained if the absence of Bbc1 enhances interaction between Myo1 and Vrp1.

Verprolins bind long-tailed myosin-1 and WASp homologs through two distinct domains, a proline-rich region and the C-terminal WASp binding domain, respectively (Anderson et al., 1998; Carnahan and Gould, 2003; Naqvi et al., 1998; Ramesh et al., 1997; Sirotkin et al., 2005). Therefore, *S. pombe* Vrp1 can bridge Myo1-Wsp1 association and may help retain both Myo1 and Wsp1 in patches. We hypothesized that increased interaction of Bbc1 with Myo1 upon Bbc1 overexpression will disrupt this bridging interaction and decrease accumulation of Vrp1, Myo1, and Wsp1 in patches (Figure 4E). Indeed, the levels of Vrp1, Myo1, and Wsp1 were significantly reduced in Bbc1 overexpressing cells (Figure 4, F-H, and Supplemental Figure S4, A and B). Collectively, these data support the competitive binding of Bbc1 and Vrp1 to Myo1 in cells, with Vrp1 promoting and Bbc1 reducing accumulation of Myo1 in patches.

### Bbc1 and Sla1 inhibit assembly of NPFs and actin at endocytic sites

Actin patch assembly is enhanced in *S. cerevisiae* cells lacking Bbc1 (Picco et al., 2018) and to even greater degree in cells lacking both Bbc1 and the adaptor protein Sla1 (Kaksonen et al., 2005). To test if inhibition of actin assembly by Sla1 and Bbc1 is conserved in *S. pombe*, we examined the effects of *bbc1*Δ alone or in combination with *sla1*Δ on accumulation in patches of F-actin labeled with LifeAct-mCherry, Crn1-GFP, or Fim1-mGFP (Huang et al., 2012; Pelham and Chang, 2001; Sirotkin et al., 2010) and NPFs Myo1, Wsp1, Vrp1, and Pan1 (Figure 5 and Supplemental Figure S5).

In agreement with previous observations (Figure 4), the loss of Bbc1 alone resulted in mild increase in accumulation of Myo1 in patches and notable increase in accumulation of Vrp1 in patches, but had no effect on accumulation of Pan1-GFP in patches (Figure 5, A-C, and Supplemental Figure 5, A and B). In contrast to the increased accumulation of WASp Las17 and actin in patches reported in *S. cerevisiae bbc1Δ* cells (Kaksonen et al., 2005; Picco et al., 2018), the peak levels and lifetimes in patches for mGFP-Wsp1 and F-actin markers Crn1-GFP and Fim1-mGFP were unaltered in *S. pombe bbc1Δ* cells (Figure 5D and Supplemental Figure S5, C-E). Likewise, even though F-actin levels observed with LifeAct-mCherry were inconsistent between experiments, the mean intensities indicated no change in actin levels in the presence or absence of Bbc1 (Figure 5E and Supplemental Figure S5F).

**Figure 5:**
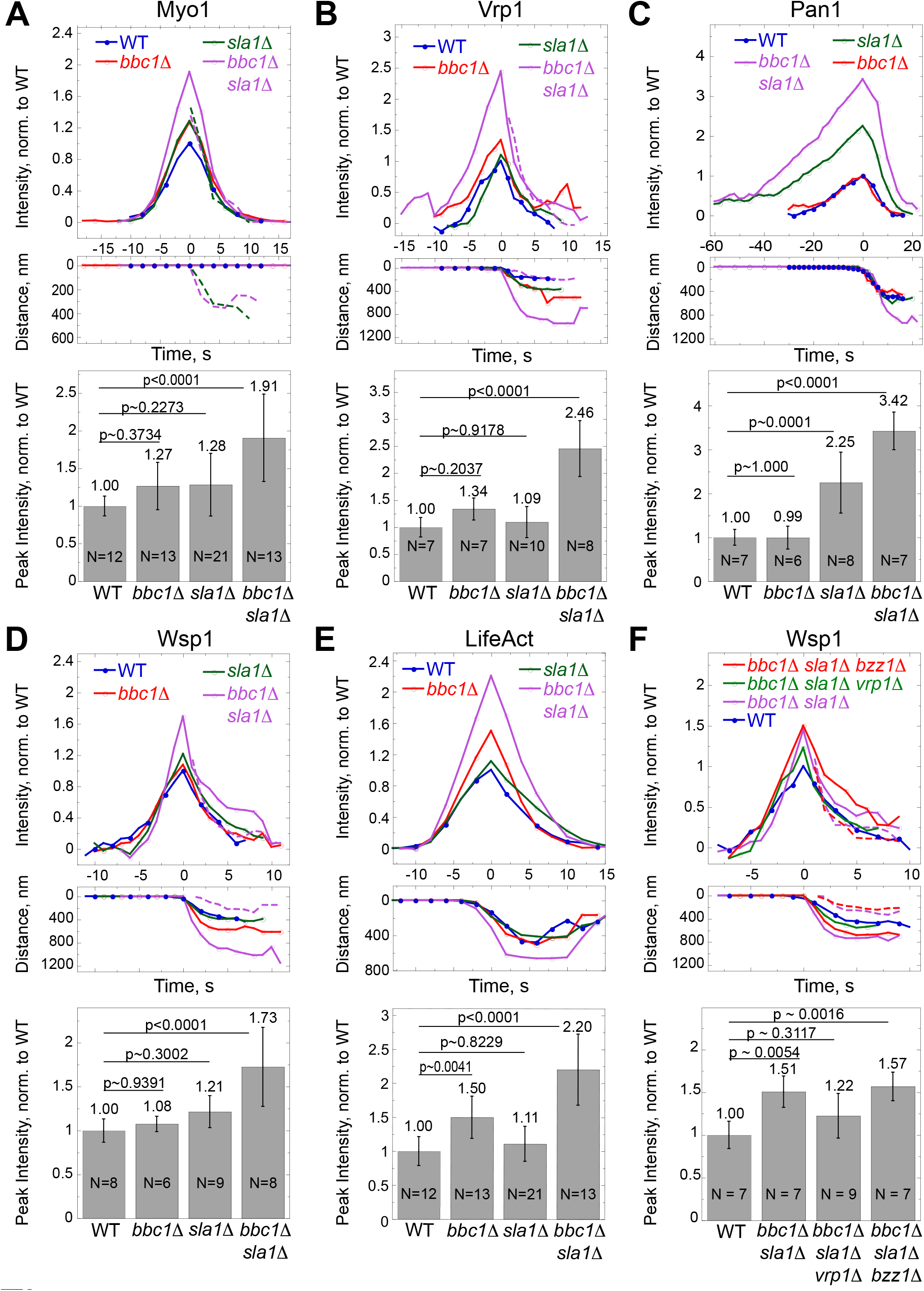
Bbc1 and Sla1 together limit accumulation of NPFs and actin in patches. (A-F) Average time courses of intensities and distances traveled, and bar graphs of average peak intensities of (A) Myo1-GFP, (B) Vrp1-mGFP, (C) Pan1-GFP, (D) mGFP-Wsp1, (E) LifeAct-mCherry in patches in WT (blue), *bbc1Δ* (red), *sla1Δ* (green), and *bbc1Δ sla1Δ* (purple) cells, and (F) mGFP-Wsp1 in *bbc1Δ sla1Δ bzz1Δ* (red), *bbc1Δ sla1 Δ vrp1 Δ* (green), *bbc1 Δ sla1 Δ* (purple), and WT (blue) cells. Dashed lines in (A-C, and F) represent the fluorescence intensities and distance traveled by portions of patches exhibiting unusual behavior: (A) internalizing portions of Myo1 patches or (B-C, and F) basal portions of splitting Wsp1 and Vrp1 patches. Time courses for individual patches were aligned to the time each patch reached peak intensity (t=0), averaged at each time point, and normalized to the average peak intensities in WT cells. Error bars represent SD. N indicates number of patches analyzed. P values represent statistical significance as determined by ANOVA with Tukey’s Post-Hoc test.

Examination of *sla1Δ* cells revealed only modest increases in accumulation of all markers except Pan1. Compared to wild-type cells, peak patch intensities of Myo1-GFP, Vrp1-mGFP, mGFP-Wsp1, and LifeAct-mCherry in *sla1Δ* cells increased by 28%, 9%, 21% and 11%, respectively (Figure 5A, B, D, and E), while the peak intensity of Pan1-GFP increased by 125% (Figure 5C). In addition, Pan1 patch lifetime was significantly increased from 35 seconds to 64 seconds. The reasons for these dramatic effects of *sla1Δ* on Pan1 are not clear but a similar increase of Pan1 patch lifetime in *sla1Δ* cells has been reported in *S. cerevisiae* (Kaksonen et al., 2005).

Combining *bbc1Δ* with *sla1Δ* resulted in greatly increased accumulation of NPFs and F-actin in patches. Compared to wild-type cells, in *sla1Δ bbc1Δ* cells, Myo1-GFP, Vrp1-mGFP, Pan1-GFP, mGFP-Wsp1, and LifeAct-mCherry peak patch intensities increased by 91%, 146%, 242%, 51-73%, and 120%, respectively (Figure 5). The same trends were observed with N-terminally tagged Myo1 and Fim1-mGFP, but to a lesser extent (Supplemental Figures S5A and S5E). Combining *vrp1Δ* but not *bzz1Δ* with *bbc1Δ sla1Δ* reduced Wsp1 accumulation, indicating that Vrp1 helps recruit or retain Wsp1 in patches (Figure 5F). The synergistic rather than additive effects of *sla1Δ* and *bbc1Δ* on assembly of NPFs and actin into patches suggest that *S. pombe* Sla1 and Bbc1 play redundant roles in constraining actin patch assembly by limiting accumulation of NPFs in patches and modulating their localization and interactions.

### Myo1 partially internalizes with actin patches in the absence of Sla1

In addition to increased accumulation of Myo1 in patches, *sla1Δ* and *bbc1Δ sla1Δ* cells displayed an unusual behavior of Myo1 at the endocytic sites. In wild-type and *bbc1Δ* cells, Myo1 always remains at the base of endocytic invaginations and does not internalize with vesicles (Sirotkin et al., 2005). In contrast, in 6% of Myo1-mGFP patches in *sla1 Δ* cells, a small portion of Myo1 moved into the cytoplasm, presumably in association with the tip of the endocytic invagination or endocytic vesicle, while a larger portion of Myo1 remained at the base (Figures 6, A and B). This partial internalization of Myo1, observed as splitting of the Myo1 signal, was significantly more pronounced in *bbc1*Δ *sla1*Δ cells where the percent of Myo1 patches exhibiting this behavior increased from 6% to 64% (Figures 6, A and B) and was observed with both N-terminally and C-terminally tagged Myo1 (data not shown). In *bbc1*Δ *sla1*Δ cells, Myo1 patches were brighter and the internalizing portion of Myo1 moved almost two times farther than in *sla1*Δ cells (Figure 6C), making Myo1 splitting events much more evident.

**Figure 6:**
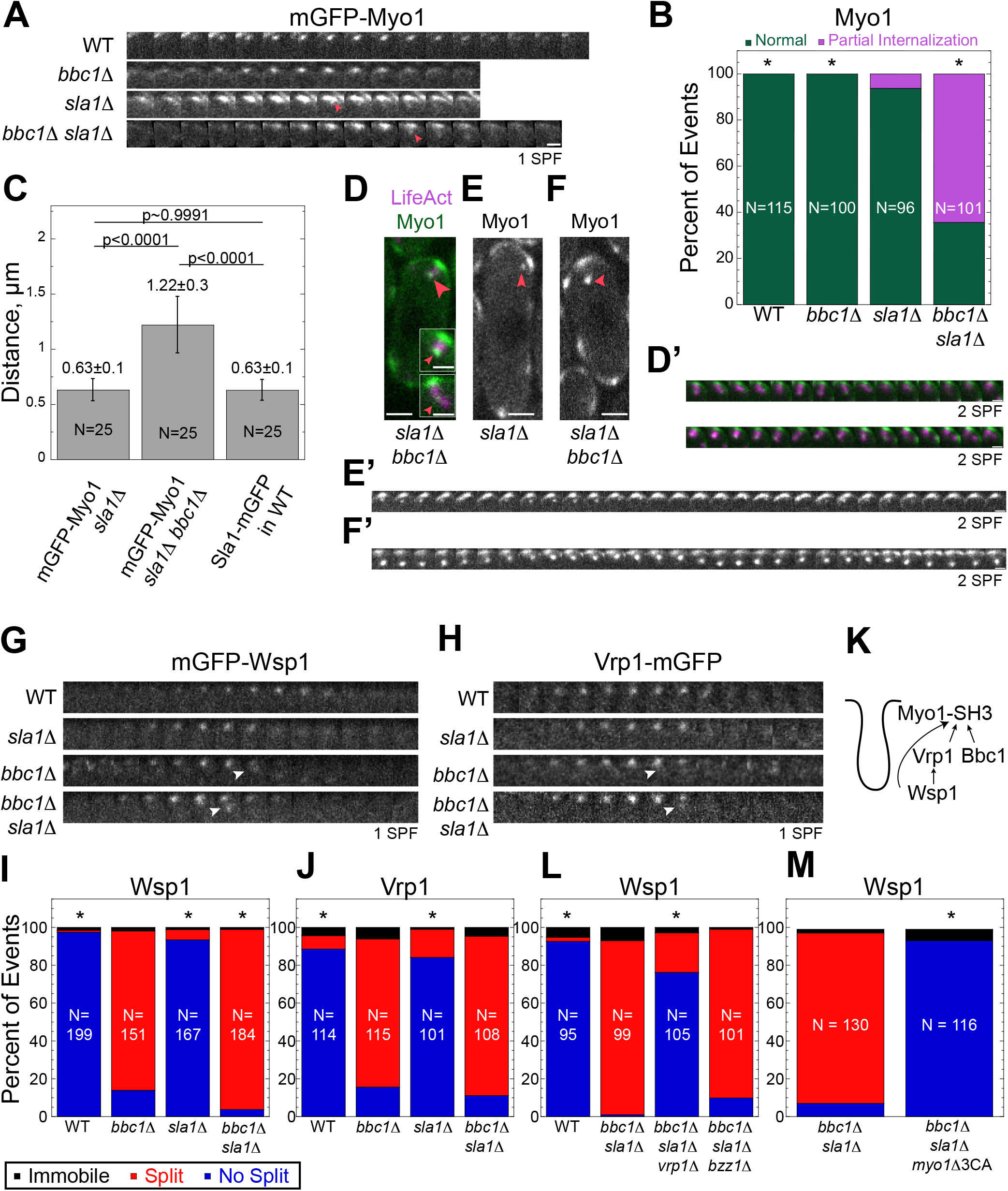
Myo1 partially internalizes in the absence of Sla1, while Wsp1 and Vrp1 patches split in the absence of Bbc1. Vrp1 and the Myo1 SH3 domain mediate Wsp1 patch splitting. (A) Time lapse montages showing mGFP-Myo1 patches partially internalizing in *sla1 Δ* and *bbc1 Δ sla1 Δ* but not in *bbc1Δ* and WT cells. Arrowheads mark instances of Myo1 patch partial internalization. Images were taken at the rate of 1 SPF. Scale bars, 1 μm. (B) Quantitation of the percent of Myo1 patches partially internalizing (magenta) and behaving normally (not internalizing, green). Asterisks indicate a significant difference from *sla1 Δ* as determined by Chi-squared test, *p<0.05. (C) Bar graphs of the distance traveled by the internalized portion of mGFP-Myo1 patches in *sla1 Δ* and *bbc1 Δ sla1 Δ* compared to Sla1-mGFP patches in WT cells. Error bars represent SD. P values represent statistical significance as determined by ANOVA with Tukey’s Post-Hoc test. (D-F) Single confocal sections through the middle of the cells and (D’-F’) corresponding time lapse montages showing (D, D’) localization of Myo1-GFP (green) at the base and the tip of the actin plume visualized with LifeAct-mCherry (magenta) in a *bbc1 A sla1 A* cell, (E, E’) mGFP-Myo1 in actin plumes in a *sla1 Δ* cell, and (F, F’) mGFP-Myo1 in actin plumes in a *bbc1 Δ sla1 Δ* cell. Insets in (D) show examples of Myo1-GFP at the tip and kink in the actin plume (arrowheads). Other arrowheads in (D-F) indicate Myo1 spots shown on montages in (D’-F’). Images were acquired at the rates of 2 SPF (D’) and 1 SPF (E’-F’) but are displayed at the intervals of 2 SPF in all panels. Scale Bars, (D-F) cells, 2 μm, (D’-E’) and insets in (D), 1 μm. (G-H) Montages of time lapse images of (G) mGFP-Wsp1 and (H) Vrp1-mGFP patches splitting in *bbc1 Δ* and *bbc1 Δ sla1 Δ* cells and not splitting in *sla1Δ* and WT cells. Scale bar, 1 μm. Arrowheads indicate instances of patch splitting. Images were taken at the intervals of 1 SPF. (I, J) Quantification of the percent of events when (I) mGPF-Wsp1 or (J) Vrp1-mGFP patches internalized normally (blue), remained immobile (black), or split in two (red) in WT, *bbc1 Δ, sla1 Δ*, and *bbc1 Δ sla1 Δ* cells. Asterisks indicate a significant difference from *bbc1 Δ* as determined by Chi-squared test, *p<0.001. (K) Diagram of proposed interactions of the Myo1 SH3 domain. (L, M) Effects of (L) *vrp1 A* or *bzz1 A* and (M) *myo1*Δ*SH3-LCA* (*myo1Δ3CA*) on the percent of events when mGPF-Wsp1 patches internalized normally (blue), remained immobile (black), or split in two (red) in *bbc1 Δ sla1 Δ* cells. Asterisks indicate a significant difference from *bbc1 Δ sla1 Δ* as determined by Chi-squared test, *p<0.001. N indicates number of patches analyzed.

We also observed infrequently in *sla1*Δ and, more often, in *sla1*Δ *bbc1*Δ cells elongated actin structures that we called actin plumes (Figures 6, D and D’ and Supplemental Video 1). These are similar to enlarged actin structures found in *S. cerevisiae sla1Δ bbc1Δ* cells (Kaksonen et al., 2005). In addition to localizing to the base of these plumes, Myo1 associated with both the tip of the actin plumes and at the middle where the plume appeared to kink (Figure 6D, inset). In single color imaging of mGFP-Myo1 in *sla1 Δ* and *bbc1 Δ sla1 Δ* cells, these plumes appeared as stable cytoplasmic dots of Myo1 swinging like a pendulum from stable Myo1 platforms on the membrane (Figures 6, E-F’ and Supplemental Video 2). Thus, besides suppressing patch assembly, Sla1 appears to play a role in regulating Myo1 positioning at the endocytic site.

### Bbc1 disrupts a transient interaction of Myo1 with Wsp1 and Vrp1

Close inspection of mGFP-Wsp1 dynamics revealed that this NPF also exhibits an unusual behavior but in *bbc1Δ* and *bbc1Δ sla1Δ* rather than *sla1 A* cells. Unlike *S. cerevisiae* Las17 and Vrp1 (Kaksonen et al., 2003; Kaksonen et al., 2005), *S. pombe* Wsp1 and Vrp1 normally internalize with an endocytic vesicle for a few seconds before dissipating into the cytoplasm (Sirotkin et al., 2005). However, in *bbc1Δ* and *bbc1Δ sla1Δ* cells, Wsp1 and Vrp1 patches frequently split in two, with a fraction internalizing normally and another fraction remaining at the base of invagination (Figures 6, G and H, and Supplemental Video 3). 84% and 95% of mGFP-Wsp1 patches split in *bbc1Δ* and *bbc1*Δ *sla1*Δ cells, respectively, while mGFP-Wsp1 patches rarely split in *sla1 Δ* and wild-type cells (Figure 6I). Vrp1-mGFP patches split at a similar frequency: 78% in *bbc1Δ*, 84% in *bbc1Δ sla1Δ* cells, but fewer than 15% of Vrp1-mGFP patches split in *sla1*Δ and wild-type cells (Figure 6J).

Since Vrp1 can bind both Myo1 and Wsp1 and thereby bridge Myo1-Wsp1 interaction (Carnahan and Gould, 2003; Sirotkin et al., 2005), splitting of Wsp1 patches in *bbc1*Δ cells could be due to enhanced association of Wsp1 at the base of the endocytic invagination with Myo1 via Vrp1 (Figure 6K), which accumulates in patches at higher levels in the absence of Bbc1 (Figure 4B). To determine if Vrp1 mediates Wsp1 patch splitting, we examined the effects of Vrp1 loss on Wsp1 patch splitting in *bbc1*Δ *sla1*Δ cells where Wsp1 patches are brighter and splitting is easier to observe (Figures 5B and 6G). Indeed, the frequency of Wsp1 patch splitting decreased from 92% in *bbc1*Δ *sla1*Δ cells to 21% in *bbc1*Δ *sla1*Δ *vrp1*Δ cells (Figure 6L). In contrast, the loss of Bzz1, which is another Wsp1 interactor that localizes to the base of endocytic invaginations (Arasada and Pollard, 2011; Idrissi et al., 2012; Kishimoto et al., 2011; Soulard et al., 2002; Sun et al., 2006), had no effect on Wsp1 patch splitting (Figures 6L) or Wsp1 accumulation in patches in *bbc1*Δ *sla1*Δ cells (Figure 5F). A reduction of Wsp1 patch splitting was also observed when *vrp1*Δ, but not *bzz1*Δ, was combined with just *bbc1*Δ (Supplemental Figure S6A), while in just *vrp1*Δ cells, as in wild-type cells, Wsp1 patches rarely split.

To test if the residual Wsp1 patch splitting observed in *bbc1*Δ *sla1*Δ *vrp1*Δ cells is due to direct binding of Wsp1 to Myo1 SH3 that was observed by two-hybrid analysis (Carnahan and Gould, 2003), we combined *bbc1*Δ *sla1*Δ with the deletion of the Myo1 SH3 and LCA domains to disrupt Myo1 interaction with both Vrp1 and Wsp1. As predicted, Wsp1 patch splitting completely ceased in *bbc1*Δ *sla1*Δ *myo1*Δ*SH3-LCA* cells (Figure 6M), suggesting that both indirect (i.e. Vrp1-mediated) and direct binding of Wsp1 to Myo1 are responsible for Wsp1 patch splitting in the absence of Bbc1. We further confirmed that Wsp1 patch splitting is mediated by the Myo1 SH3 domain as deletion of just the Myo1 CA domain had only a minor but statistically significant effect on splitting of Wsp1 patches in *bbc1*Δ and *bbc1*Δ *sla1*Δ cells (Supplemental Figure S6B).

### Bbc1 Regulates Endocytic Invagination Length

The observation that Myo1-mGFP patches internalized nearly twice as far in *bbc1Δ sla1Δ* cells compared to *sla1Δ* cells (Figure 6C) suggested that, similar to *S. cerevisiae* Bbc1 (Kaksonen et al., 2005; Picco et al., 2018), *S. pombe* Bbc1 might have a role in regulating the length of the endocytic invagination. To determine the effect of *bbc1*Δ on the length of endocytic invaginations in *S. pombe* (Figure 7A), we measured the accumulation of amphyphisin Hob1, whose *S. cerevisiae* homolog Rvs167 binds around the tubular invagination (Idrissi et al., 2012; Idrissi et al., 2008; Kishimoto et al., 2011; Kukulski et al., 2012; Picco et al., 2015), and the distances traveled by Hip1R homolog End4/Sla2 and epsin Ent1 that in *S. cerevisiae* associate with the tips of endocytic invaginations (Boettner et al., 2011; Idrissi et al., 2012; Picco et al., 2015; Skruzny et al., 2012). Compared to wild-type cells, Hob1-mGFP patch accumulation increased by 88% in *bbc1Δ* cells and 77% in *bbc1Δ sla1Δ* cells but it did not increase in *sla1Δ* cells (Figure 7, B and C). Additionally, the average distance traveled by Hob1-mGFP patches doubled in both *bbc1Δ* and *bbc1Δ sla1Δ* mutants compared to patches in wild-type and *sla1Δ* cells (Figure 7C). Similarly, End4-mGFP and Ent1-mGFP patches internalized approximately 2 times farther in *bbc1Δ* and *bbc1Δ sla1Δ* cells than in wild-type and *sla1Δ* cells: ~1 μm in *bbc1Δ* and *bbc1Δ sla1Δ* cells versus ~0.5 μm in wild-type and *sla1Δ* cells (Figures 7, D and E). The increased distance measured for End4-mGFP patches is not a result of the ability to track brighter patches longer since End4-mGFP patches had similar intensities in wild-type and *bbc1Δ* cells (Supplemental Figure S7A). Likewise, another invagination tip marker, Sla1 (Idrissi et al., 2008; Kukulski et al., 2012), moved twice as far in *bbc1Δ* cells compared to wild-type cells (Supplemental Figure S7B). Collectively, these measurements consistently reveal a 2-fold increase in the length of endocytic invaginations in cells lacking Bbc1, which we hypothesized was due to prolonged association of Myo1 with Vrp1 and Wsp1, the main NPF at the endocytic site (Berro et al., 2010), at the base of endocytic invaginations. In agreement with this hypothesis, the distance traveled by the tip marker End4-mGFP returned to normal 0.5 μm when *bbc1Δ* or *bbc1Δ sla1Δ* were combined with *myo1ΔSH3-LCA*, thereby preventing Wsp1 retention at the base of invagination (Figure 7F).

**Figure 7:**
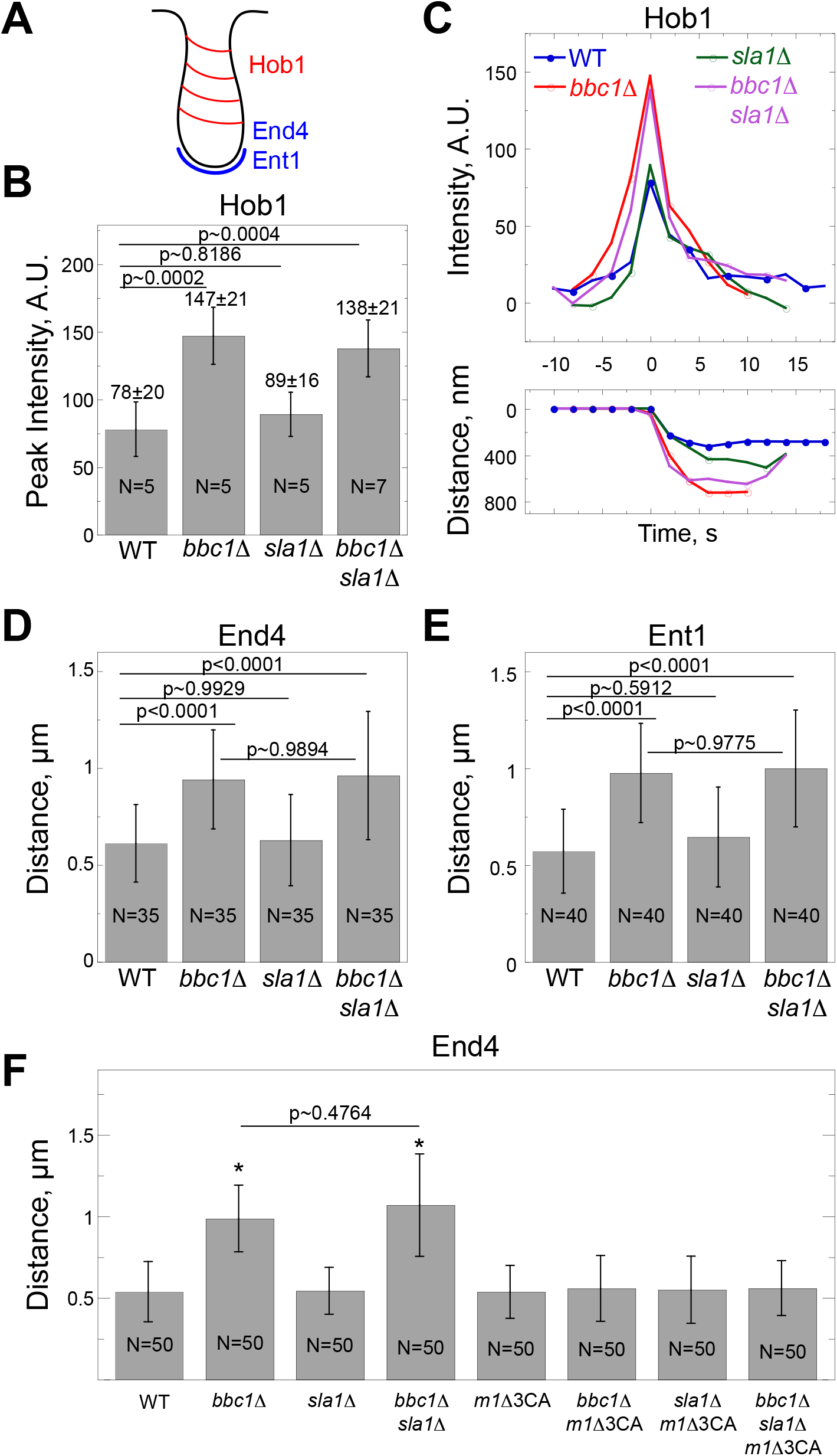
Bbc1 regulates endocytic invagination length. (A) A schematic of the proposed localization of Hob1, End4, and Ent1 at the endocytic site. (B) Bar graph showing average peak intensity of Hob1-mGFP in patches and (C) average time courses of intensities and distances traveled for Hob1-mGFP in patches in WT (blue), *bbc1Δ* (red), *sla1Δ* (green), and *bbc1Δ sla1Δ* (purple) cells. Time courses for individual patches were aligned to the time each patch reached peak intensity (t=0), averaged at each time point, and normalized to the average peak intensities in the WT cells. (D, E) Distances traveled by (D) End4-mGFP and (E) Ent1-mGFP in endocytic patches in WT, *bbc1Δ, sla1Δ*, and *bbc1Δ sla1Δ* cells. (F) Effect of *myo1ΔSH3-LCA (m1Δ3A)* on distance traveled by End4-mGFP in cells with different combinations of *bbc1Δ* and *sla1Δ*. Error bars represent SD. N indicates number of patches analyzed. P values in all panels represent statistical significance as determined by ANOVA with Tukey’s Post-Hoc test. *p<0.0001.

## DISCUSSION

In this work, we found that the previously uncharacterized *S. pombe* protein Bbc1 localizes to sites of endocytosis where it regulates NPF localization and invagination. Bbc1 is recruited to actin patches in a Myo1-dependent manner where it co-localizes with the F-BAR protein Cdc15. In *bbc1 A* cells, Vrp1 patch accumulation is increased and Vrp1-Wsp1 atypically localize to the base of endocytic invaginations. Additionally, *bbc1Δ* cells show increased length of endocytic invaginations. Combining *bbc1Δ* with *sla1Δ* resulted in exaggerated accumulation of NPFs and actin in patches, suggesting Bbc1 and Sla1 collaborate to negatively regulate endocytosis. We propose that Bbc1 negatively regulates endocytosis not through direct inhibition of the primary NPF Wsp1, but by regulating its localization to the proper site on the endocytic tube and thus in turn promoting actin polymerization at the appropriate place.

### Bbc1 is part of an interconnected actin patch protein network

Despite Bbc1 identification in pull-down assays with CR proteins Cyk3 and Fic1, our data indicate that Bbc1 is not a CR component. Instead, Bbc1 localizes to dynamic actin patches surrounding the ring, with timing similar to other patch components (Wu et al., 2006) and consistent with Bbc1 co-purifying with actin patch proteins Pan1, Myo1, and Cdc15. Upon completion of ring constriction, in addition to actin patches, Bbc1 localizes to actin-free stable puncta at the new cell tips. Similar puncta have been reported for Cyk3, Fic1, and Cdc15 (Carnahan and Gould, 2003; Pollard et al., 2012; Roberts-Galbraith et al., 2009), suggesting that Bbc1 co-localization with these CR proteins in stable puncta may account for their co-purification.

We have established that Bbc1 is recruited to actin patches in a Myo1-dependent manner. Myo1 is also required to recruit Cdc15 (Carnahan and Gould, 2003) and interacts with Vrp1 (Sirotkin et al., 2005). Myo1’s multi-domain structure facilitates these interactions: Cdc15 interacts with the TH1 domain (Carnahan and Gould, 2003) and partially depends on TH2 and SH3 domains, while Bbc1 requires the TH2 and SH3 domains for patch localization. Indeed the Bbc1 SH3 domain could bind the proline-rich Myo1 TH2 domain, while the Myo1 SH3 domain could bind the proline-rich regions of Bbc1 simultaneously.

Vrp1 however can also bind the Myo1 SH3 domain (Sirotkin et al., 2005). Our data revealed that the absence of Vrp1 increases Bbc1 patch localization by ~40%. Reciprocally, Bbc1 loss leads to ~50% increase in Vrp1 patch accumulation. This leads to the hypothesis that Bbc1 and Vrp1 compete for binding the Myo1 SH3 domain (Figure 8).

**Figure 8:**
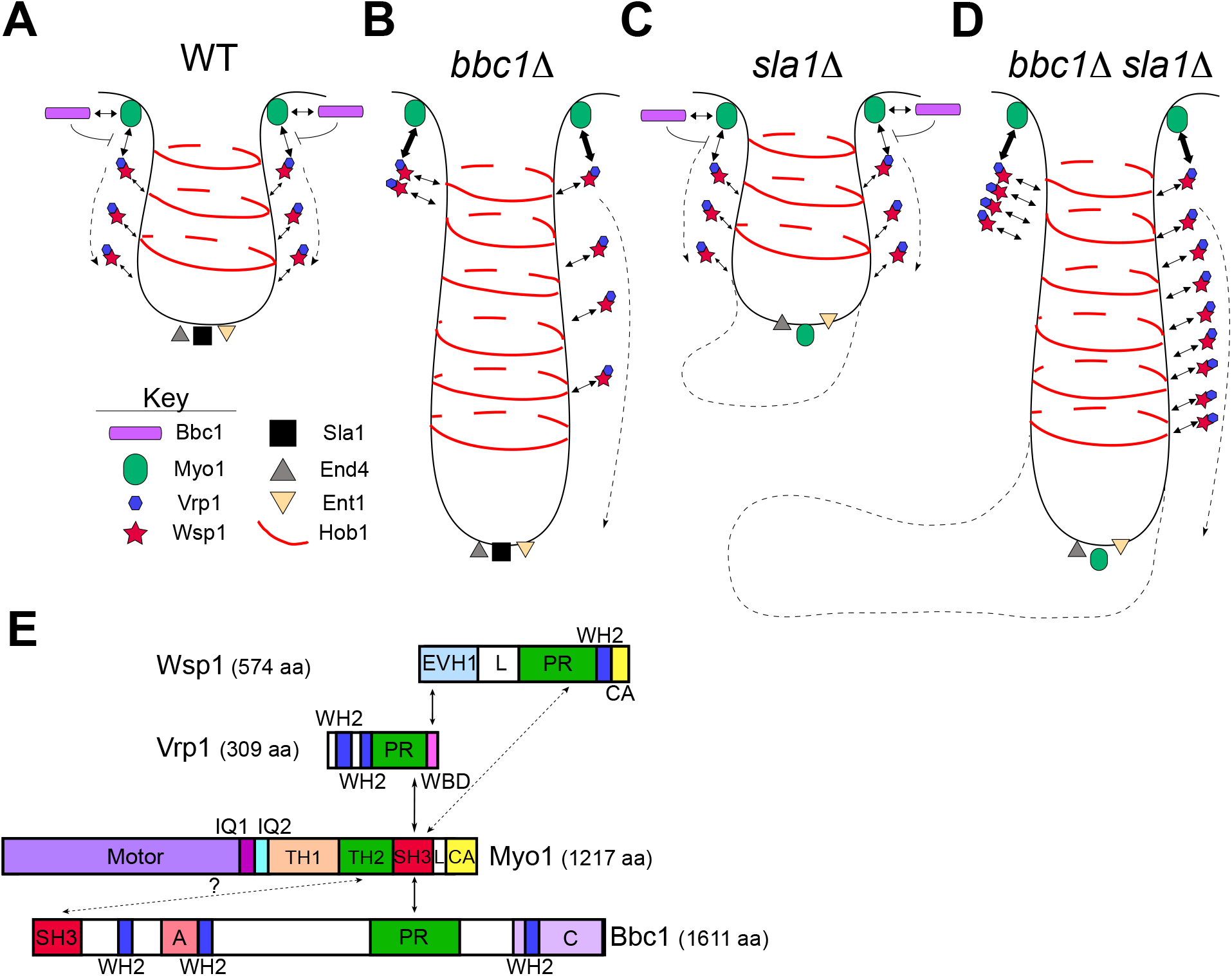
A model for regulation of actin patch assembly by Bbc1. (A-D) A graphic summary of the effects of *bbc1Δ* and *sla1Δ* on the distribution and interactions of selected endocytic patch proteins and the endocytic site organization compared to WT. (A) In WT cells, Bbc1 inhibits transient interaction of Wsp1 and Vrp1 with Myo1 at the base of invagination so that Wsp1 and Vrp1 internalize with the actin patch while Myo1 and Bbc1 remain at the base. (B) In the absence of Bbc1 (*bbc1Δ*), interactions of Wsp1 and Vrp1 with Myo1 are enhanced, resulting in retention of a portion of Wsp1 and Vrp1 at the base of the invagination and longer endocytic invaginations. (C) In the absence of Sla1 (*sla1Δ*), a portion of Myo1 associates with the tip of the invagination. (D) In the absence of both Bbc1 and Sla1, the effects of *bbc1Δ* alone are amplified by increased accumulation of NPFs and enhanced actin assembly. (E) A schematic diagram of Wsp1, Vrp1, Myo1, and Bbc1 protein domains and their known (solid lines) or proposed (dashed lines) binding interactions. EVH1, Ena/VASP homology; L, Linker, PR, Proline-Rich, WH2, WASp Homology 2; CA, Central-Acidic; WBD, WASp binding domain, TH1 and 2, Tail Homology 1 and 2, SH3, Src Homology 3, A, Glu/Asp-rich, C-term, conserved C-terminal domain.

Unlike Bbc1, which completely depends on Myo1 for localization to patches, Vrp1 is only partially dependent on Myo1 and instead requires Wsp1 for patch localization (Sirotkin et al., 2005). Consistent with our hypothesis, the absence of Wsp1 also leads to an increase in Bbc1 patch localization, presumably by loss of Vrp1 from actin patches.

Although *wsp1*Δ increases Bbc1 patch localization, *S. pombe bbc1*Δ does not alter the amount of Wsp1 in patches, nor does it affect actin abundance. Surprisingly, the major phenotype observed in *bbc1*Δ cells was the splitting of Wsp1 patches: Wsp1 normally internalizes with the endocytic tube but in *bbc1*Δ we observed some Wsp1 remaining associated with the base of the invagination.

Again, this data is consistent with a model (Figure 8) in which Bbc1 and Vrp1 compete for binding the Myo1 SH3 domain. In this case, Bbc1 loss permits increased association of Myo1 SH3 domain with Vrp1 and its binding partner Wsp1. Furthermore, this model predicts that Wsp1 patch splitting would be abrogated in *vrp1*Δ or *myo1*Δ*SH3-LCA* cells, which we also observed.

### Comparison with budding yeast

In *S. cerevisiae bbc1*Δ cells, as in *S. pombe*, patches of WASp homolog Las17 split and endocytic vesicles internalize deeper (Kaksonen et al., 2005). However, we expect that the mechanism responsible for splitting of Las17 patches in *S. cerevisiae* is likely different from that in *S. pombe*. In *S. cerevisiae bbc1*Δ cells, more Las17 accumulates at endocytic sites over a larger membrane surface and the internalized membrane area is greater than in wild-type cells (Picco et al., 2018). We suspect that splitting of Las17 patches is due to the increased internalization of Las17-coated membrane areas that normally are not internalized. Indeed, a recent study revealed that Las17 localizes in a donut-like pattern surrounding the area containing proteins found at the tip of endocytic invagination (Mund et al., Nov. 15, 2017). Interestingly, in *bbc1*Δ cells, Las17 localization spreads to a wider area containing myosin-1 Myo5. This might be due to increased association of Las17 with Myo5 in the absence of competition from Bbc1 for the myosin-1 SH3 domain, similar to the mechanism we propose for *S. pombe* Wsp1. Alternatively, *S. cerevisiae* Bbc1 might bind Las17 with appreciable affinity and this interaction may help retain Las17 at the base of the invagination. The loss of this interaction would result in internalization of a portion of Las17 observed as patch splitting.

In *S. cerevisiae*, loss of Bbc1 is thought to not only affect Las17 accumulation and localization at patches, but also to inhibit its NPF activity. *In vitro* assays demonstrated that *S. cerevisiae* Bbc1 binds WASp homolog Las17 and inhibits its NPF activity (Rodal et al., 2003; Tong et al., 2002). *In vivo* Bbc1 and Las17 both remain at the base of endocytic invaginations where presumably Bbc1 directly inhibits Las17. Several lines of evidence argue against *S. pombe* Bbc1 interacting with Wsp1 to inhibit its activity. First, unlike Myo1, Wsp1 was not detected in our pull-down assays with Bbc1. Second, Bbc1 does not depend on Wsp1 for localization to patches. Third, Bbc1 exhibits Myo1-like behavior and remains stationary at the base of the invagination, while Wsp1 and Vrp1 move away with the actin patch (Sirotkin et al., 2005). Fourth, our molecular counts show ~1:3 stoichiometry of Bbc1:Wsp1 molecules in patches (Sirotkin et al., 2010); insufficient to inhibit all of the Wsp1 in a patch. Therefore, we propose an alternative model for how Bbc1 may regulate Wsp1 by regulating its location along the endocytic invagination and proximity to activators such as Bzz1 (Arasada and Pollard, 2011).

### Bbc1 and Sla1 regulate NPF abundance and actin polymerization

In *S. cerevisiae*, Bbc1 and Sla1 both inhibit Las17 *in vitro* (Rodal et al., 2003) and *bbc1*Δ *sla1*Δ cells show enhanced actin patch assembly, sometimes resulting in the formation of actin comet tails (Kaksonen et al., 2005). While our data suggest that, as discussed above, Bbc1 may not be directly inhibiting Wsp1 NPF activity, our analysis of actin patch assembly in *bbc1*Δ *sla1*Δ cells does not rule out the proposed role for Bbc1 and Sla1 as redundant NPF inhibitors. However, our data reveals additional mechanistic insights into Bbc1 and Sla1 functions and raises the possibility that Bbc1 and Sla1 play distinct roles in regulation of NPF localization and activity (Figure 8). In *S. pombe*, either *bbc1*Δ or *sla1*Δ alone had no or only minor effects on the extent of actin patch assembly but together resulted in a 2-fold increase in actin assembly, consistent with NPF inhibition by Bbc1 and Sla1. However, this increase of actin patch assembly in *bbc1*Δ *sla1*Δ cells was accompanied by ~2-3-fold increased accumulation of all NPFs, suggesting that enhanced actin assembly is the result of a greater concentration of NPFs in patches rather than loss of inhibition of NPF activity. Moreover, *bbc1*Δ and *sla1*Δ had distinct effects on distribution of NPFs at endocytic sites.

We observed two intriguing effects of *sla1*Δ on actin patch assembly. First, in *sla1 Δ* cells, 2-fold more Pan1 assembled into patches over a longer period of time, possibly due to delayed cargo accumulation. Second, in *sla1*Δ, and more prominently in *bbc1*Δ *sla1*Δ cells, a portion of Myo1 internalized with the patch. We considered a possibility that this could be the consequence of the increased amounts of Pan1 in patches, based on the interaction of myosin-1 with Pan1 proline-rich domain (PRD) reported in budding yeast (Barker et al., 2007). However, deletion of the Pan1 PRD had no effect on Myo1 internalization in *S. pombe bbc1*Δ *sla1*Δ cells (our unpublished data). Alternatively, the SH3 domains of Myo1 and Sla1 can be competing for binding the Wsp1 PRD, so that in the absence of Sla1, normally present at the invagination tip, Myo1 binds and internalizes with Wsp1.

### Models of force generation at endocytic sites

In *S. pombe* as in other organisms, actin polymerization provides the force for overcoming membrane tension during endocytosis (Basu et al., 2014). In wild-type cells, prior to initiation of an endocytic invagination, Wsp1 and Myo1 co-localize at endocytic patches where they interact directly and indirectly via Vrp1 (Carnahan and Gould, 2003; Sirotkin et al., 2005). Vrp1 activates Myo1 NPF activity (Sirotkin et al., 2005) while Bzz1 stimulates Wsp1 NPF activity (Arasada and Pollard, 2011). As the invagination forms, Wsp1 and Vrp1 internalize while Myo1 and Bzz1 remain on the membrane at the base of the invagination (Sirotkin et al., 2005). One model of force generation suggests that this separation results in two separate zones of branched actin assembly that push off of each other (Arasada and Pollard, 2011; Arasada et al., 2018).

An alternative model proposes that most of the force is generated in one zone at the base of the emerging invagination (Picco et al., 2015). Several lines of evidence favor this latter model. First, Wsp1 is a much stronger NPF than Myo1, which also needs stimulation by Vrp1 (Sirotkin et al., 2005). Second, Wsp1 reaches peak accumulation 2 seconds before the onset of invagination, when Wsp1 and Myo1 still co-localize in the same zone (Sirotkin et al., 2010). Further, movement of Wsp1-Vrp1 with the patch would separate Wsp1 and Myo1 from their respective activators, Bzz1 and Vrp1.

In addition to splitting of Wsp1 and Vrp1 patches, *bbc1*Δ cells also have longer endocytic invaginations. We hypothesize that the prolonged localization of Wsp1 and Vrp1 at the base of the invagination leads to continuing stimulation of Wsp1 by Bzz1, further stimulation of Myo1 NPF activity by Vrp1, and orientation of the growing actin filament ends towards the plasma membrane and near Myo1, where Myo1 can exert force driving membrane invagination. We propose that this membrane-directed actin assembly is ultimately responsible for longer invaginations and increased propulsion of endocytic vesicles observed in *bbc1*Δ cells.

In sum, we have provided evidence that SH3 domains of endocytic proteins compete for their ligands and this competition plays an important role in regulating actin assembly. These observations underscore the importance of future work towards dissecting the complex network of SH3 domain interactions at endocytic sites (Fernandez-Ballester et al., 2009; Tong et al., 2002; Tonikian et al., 2009) and the role of these interactions in orchestrating the precise assembly of proteins driving endocytic internalization.

## MATERIALS AND METHODS

### Yeast strain construction

*S. pombe* strains used in this study are listed in Supplementary Table S2. LifeAct-mCherry was originally from Huang et al. (2012). All other yeast strains were constructed by PCR-based gene tagging and genetic crosses. Deletions or fluorescent protein tags marked with selectable markers were introduced into endogenous chromosomal loci by homologous recombination of PCR-amplified gene tagging modules (Bahler et al., 1998; Wach et al., 1994). Amplified modules were transformed into cells by lithium acetate method (Keeney and Boeke, 1994) and successful integrants were isolated based on selectable markers and confirmed by PCR and sequencing.

To tag proteins at the C-termini with fluorescent protein tags, the stop codons of the targeted genes were replaced with gene tagging cassettes PCR-amplified from pFA6a-mGFP-kanMX6 (pJQW 85-4) for monomeric (m) GFP (Sirotkin et al., 2010; Wu et al., 2003), pFA6a-mCherry-kanMX6 (pKS390) or pFA6a-mCherry-natMX6 (pKS391) for mCherry (Snaith et al., 2005), and pFA6a-mNeonGreen-kanMX6 for mNeonGreen (Shaner et al., 2013; Willet et al., 2015). All C-terminally tagged proteins were expressed from endogenous loci under control of the native promoters.

Gene deletions were constructed by replacing gene open reading frames with *ura4*^+^ cassette amplified from KS-ura4 (Bahler et al., 1998) for *bbc1*Δ::*ura4*^+^ and *vrp1*Δ::*ura4*^+^, *kanMX6* cassette amplified from pFA6a-kanMX6 (Bahler et al., 1998) for *bbc1*Δ::*kanMX6* and *bzz1*Δ::*kanMX6*, and *natMX6* cassette amplified from pCR2.1-nat (Sato et al., 2005) for *sla1*Δ::*natMX6*.

The C-terminal truncations of Myo1 were constructed by replacing sequences encoding domains targeted for deletion with a stop codon followed by the *ura4*^+^ cassette amplified from KS-ura4 (Bahler et al., 1998). The unmarked *myo1*Δ*TH2* allele was constructed by replacing *ura4*^+^ cassette of *myo1*Δ*TH2-SH3-LCA* allele with the SH3-LCA-stop codon sequence amplified by PCR with primers flanked by 70 nt regions of homology to the end of TH1 domain and *myo1*^+^ 3’ UTR. The *myo1*Δ*TH2* transformants were isolated by counterselection on EMM plates containing 2 mg/ml 5-Fluoroorotic Acid 9 (5-FOA) (Stark et al., 2013). The Myo1 domain boundaries were defined as previously described (Lee et al., 2000; Sirotkin et al., 2005): TH2, aa 967-1112; SH3, aa 1113-1163; L, aa 1164-1186; and CA, aa 1187-1217.

A gene expressing N-terminally tagged mCherry-Cdc15 from the native locus under control of endogenous *Pcdc15* promoter was constructed by replacing *ura4*^+^ cassette in a *cdc15*^+^/*cdc15*Δ::*ura4*^+^ diploid strain with the cassette consisting of 500 bp 5’ *cdc15*^+^ flanking region, mCherry coding sequence, a GGGGSGGGGSG linker sequence, *cdc15*^+^ coding sequence, and 500 bp 3’ *cdc15*^+^ flanking region. Haploid mCherry-Cdc15 integrants were isolated by 5-FOA selection and confirmed by sequencing.

Strains combining two or more deletion or tagged alleles were constructed by standard genetic crosses on ME (Malt Extract) plates and tetrad dissections on YES plates followed by screening for wanted gene combinations by replica plating onto appropriate selective plates, microscopy, and PCR diagnostics. In some cases, *myo1*Δ and *wsp1*Δ cells were transformed with pUR-myo1^+^ and pUR-wsp1^+^ plasmids (Lee et al., 2000), respectively, which facilitate mating and are subsequently lost upon sporulation.

### Plasmid construction

Ppc89-Linker-mCherry cassette flanked by XhoI and NheI sites in pCR-BluntII-TOPO vector (Invitrogene) was created by In-Fusion cloning using ClonTech In-Fusion HD Cloning Kit (Takara Bio USA) following manufacturer’s instructions. Briefly, a linear fragment encoding a start codon (bolded), aa 261-783 of Ppc89, and a 5 aa GSGSG Linker (italicized) was PCR-amplified from pREP81-ppc89 f5/r3 with the primers Ppc89F (<u>CCTTCACTCGAG**ATG**</u>ACAATAGATGAAAACGGTAATGC) and Ppc89R (<u>CTCGCCCTTGCTCAC</u>*ACCAGAACCAGAACC*GCTAAAATCCTTACCAACGTTG). The resulting Ppc89-Linker product was fused by recombination via the 15 nt homology regions (underlined in primer sequences) to the N-terminal sequence of previously cloned mCherry in pCR-BluntII-TOPO vector that was linearized by PCR amplification with the TmChF primer (<u>GTGAGGAAGGGGGAG</u>GAG) corresponding to mCherry aa 2-7 and TmChR primer (<u>CATCTCGAGTGAAGG</u>GCGA) corresponding to the start codon, XhoI site, and pCR-BluntII-TOPO MCS. The resulting Ppc89-Linker-mCherry fragment flanked by XhoI and NheI sites was cleaved from pCR-BluntII-TOPO vector and cloned in place of mGFP in the previously made pSGP573-3xPnmt1-mGFP-L-23A and −3A constructs (Siam et al., 2004) digested with XhoI and NheI. In the resulting constructs, sequences encoding Myo1 TH2-SH3-LCA (23A) fragment (aa 967-1217) or SH3-LCA (3A) fragment (aa 113-1217) are fused to the C-terminus of Ppc89-Linker-mCherry via a 7-aa linker with the sequence GASGTGS, and the expression is controlled by the thiamine-repressible 3xPnmt1 promoter (Maundrell, 1990).

Untagged Bbc1 overexpression constructs were made by subcloning *bbc1*^+^ ORF into the XhoI and XmaI sites of pREP3x vector (Forsburg, 1993). To N-terminally tag Bbc1 with mEGFP, the Bbc1 overexpression construct was linearized with XhoI and a fragment encoding mEGFP followed by GGGGSGGGGSG linker was fused in through Gibson cloning (Gibson et al., 2009).

All plasmid constructs were verified by sequencing and then electroporated into cells followed by selection on appropriate EMM plates containing 5 μg/ml thiamine and lacking uracil for pSGP573 constructs or leucine for pREP3x constructs.

### Growth assays

Cells were grown to log phase at 25°C in YES medium. Forty million cells were resuspended in 1 ml YES and then serially diluted 10-fold four times. Two μl of each dilution were spotted to YES plates and, for drug sensitivity assays, YES plates containing 0.25 μM latrunculin A (Cayman Chemical) or 10 μg/ml benomyl (Sigma). Plates were incubated at different temperatures as indicated on the Figure S1.

### Microscopy

Except for Figures 1A-D, 2B, 2C, and S1C, imaging was performed on a UltraView VoX Spinning Disc Confocal System (PerkinElmer) mounted on a Nikon Eclipse Ti-E microscope with a 100x/1.4 NA PlanApo objective, equipped with a Hamamatsu C9100-50 EMCCD camera, and controlled by Volocity (PerkinElmer) software. Stably integrated *S. pombe* strains were grown to exponential phase (OD_595_=0.2-1.0) in liquid YES medium (Sunrise) while shaking in the dark at 25° over 2 days. Cells containing protein expression plasmids with *3xPnmt1* promoter were grown overnight at 25° in liquid EMM medium (Sunrise) lacking uracil (for pSGP573 constructs) or leucine (for pREP3x constructs) and containing 5 μg/ml thiamine to suppress premature protein overexpression, washed next day 3 times with EMM lacking thiamine, and grown overnight at 25° for 15-20 hours in the absence of thiamine to induce protein expression in liquid EMM medium lacking uracil (for pSGP573 constructs) or leucine (for pREP3x constructs). For microscopy, cells from 0.5-1 ml of culture were collected by a brief 1-minute centrifugation in a microfuge (6,000 rpm), washed with EMM, pelleted, and 5 μl of partly re-suspended pellet was placed onto a pad of 25% gelatin in EMM on a glass slide, covered by a coverslip and sealed with VALAP (1:1:1 Vaseline:Lanolin:Paraffin). Samples were imaged after a 5-minute incubation to allow for partial depolarization of actin patches. For time-lapse imaging, single color images were taken every second for 60-120 seconds and two-color images were taken every 2 seconds for 60-120 seconds. Z-series of images spanning the entire cell width were captured at 0.4 μm intervals.

Images in Figures 1A-B were acquired using a personal DeltaVision microscope system (GE Healthcare, Issaquah, WA), which includes an Olympus IX71 microscope, 60X NA 1.42 PlanApo and 100X NA 1.40 UPlanSApo objectives, fixed- and live-cell filter wheels, a Photometrics CoolSnap HQ2 camera, and softWoRx imaging software. Images were acquired at 25-29°C. Images are maximum-intensity projections of z sections spaced at 0.5 μm that have been deconvolved with 10 iterations. For images in Figure 1A, 1 ml of log-phase cells grown overnight in YES at 25°C in a shaking water bath were collected by centrifugation, resuspended in ~10 μl YES, and spotted directly on glass slide. Time-lapse imaging in Figure 1B was performed using an ONIX microfluidics perfusion system (CellASIC, Hayward, CA). Cells were grown and collected as for Figure 1A except that 40 millions cells were resuspended in 1 ml YES. Then, 50 μl cell suspension were loaded into Y04C plates for 5 s at 8 psi, and YES liquid medium flowed into the chamber at 5 psi throughout imaging. Time-lapse images were obtained at 4-minute intervals.

Images in Figures 1D, 2B and 2C were acquired with a spinning disk system, which includes a Zeiss Axiovert200m microscope, Yokogawa CSU-22, 63X NA 1.46 planApochromat and 100X numerical aperture (NA) 1.40 PlanApo oil immersion objectives, and Hamamatsu ImageEM-X2 camera. For end-on imaging, cells grown to log-phase in EMM plus adenine, uracil and leucine were loaded to vertical chambers in a 4% MAUL agarose pad and imaged every 2 minutes. To image patches in Figure 2B and 2C, cells were grown to log-phase at 25°C in MAUL and then mounted to 2% MAUL agarose pad for imaging.

For visualization of nuclei and septa in Figure S1C, cells were fixed with ice-cold 70% ethanol and incubated at 4°C for at least 30 minutes. After washing with potassium-free PBS, fixed cells were incubated in 1 mg/ml methyl blue (MB) for 30 min at room temperature (RT). MB-stained cells were pelleted by centrifugation, resuspended in PBS, and mixed with DAPI in 1:1 ratio immediately prior to imaging.

### Image Analysis

Image analysis was performed in ImageJ (National Institutes of Health). To make sure that different mutant backgrounds did not alter the expression levels of tagged proteins and that imaging conditions remain stable throughout an imaging session, for all strains within each dataset, we measured average background-subtracted whole cell intensities, which correspond to the tagged protein expression levels. For each time series, we measured whole cell intensities of 5 cells, subtracted these values for either extracellular background or the intensities of untagged wild-type cells, and averaged the background-subtracted values at each time point. All strains within each dataset had statistically similar whole cell intensities.

The intensities and positions of fluorescent protein-tagged markers in individual endocytic actin patches were manually tracked throughout lifetime of each patch using a circular ROI with a 10-pixel diameter. Mean fluorescence intensities of patches were subtracted for mean cytoplasmic intensities measured in cell areas away from patches, and distances traveled by patches from the original positions were calculated. Except for measurements of the number of molecules, time courses of background-subtracted intensities and distances from origin for individual patches were aligned to the time of peak intensity (time = 0) and averaged at each time point. Where noted, fluorescence intensities for each condition were normalized to the average peak intensity in the wild-type (WT) cells. The statistical significance of the data was tested by Student’s t-test in Microsoft Excel for datasets of only two conditions and by ANOVA with Tukey’s Post-Hoc test for datasets of more than two conditions. The percent of patches that were splitting were measured by manually following the fates of at least 80 patches for each condition and the statistical significance of the data was tested by chi-squared analysis in Excel.

For the measurements of the numbers of molecules, strains combing Fim1-mCherry and one of mGFP-tagged patch markers were imaged in a single medial plane at the rate of 1 frame every 2 seconds. Whole cell intensities and the intensities and positions of patches were measured as described above. For these measurements, the time courses of cytoplasmic background-subtracted intensities and distances from origin for individual patches were aligned to the time Fim1-mCherry patch started moving (time = 0), corresponding to the initiation of patch internalization. The intensities and distance values were averaged at each time point and the fluorescence intensities were converted into the numbers of molecules using the calibration curve constructed based on the previously measured average peak numbers of molecules of End4, Pan1, Wsp1, and Myo1 in patches, reported in Figure 7 in Sirotkin et al. (2010).

### Mass-spectrometry

Mass-spectrometry analysis was performed as previously described in Roberts-Galbraith et al. (2009; 2010).

## ACKNOWLEDGEMENTS

We thank Ms. Keely Macmillan for help with making Myo1 C-terminal truncations and Ms. Elisabeth Barone for help with making strains and tracking patches for counting molecules. This work was supported by Public Health Service awards from the NIGMS T32 GM007347 to the Vanderbilt MSTP, F31 GM119252 to M.M., and R01 GM101035 to K.L.G.

## SUPPLEMENTAL MATERIALS

### SUPPLEMENTAL FIGURE LEGENDS

**FIGURE S1:**
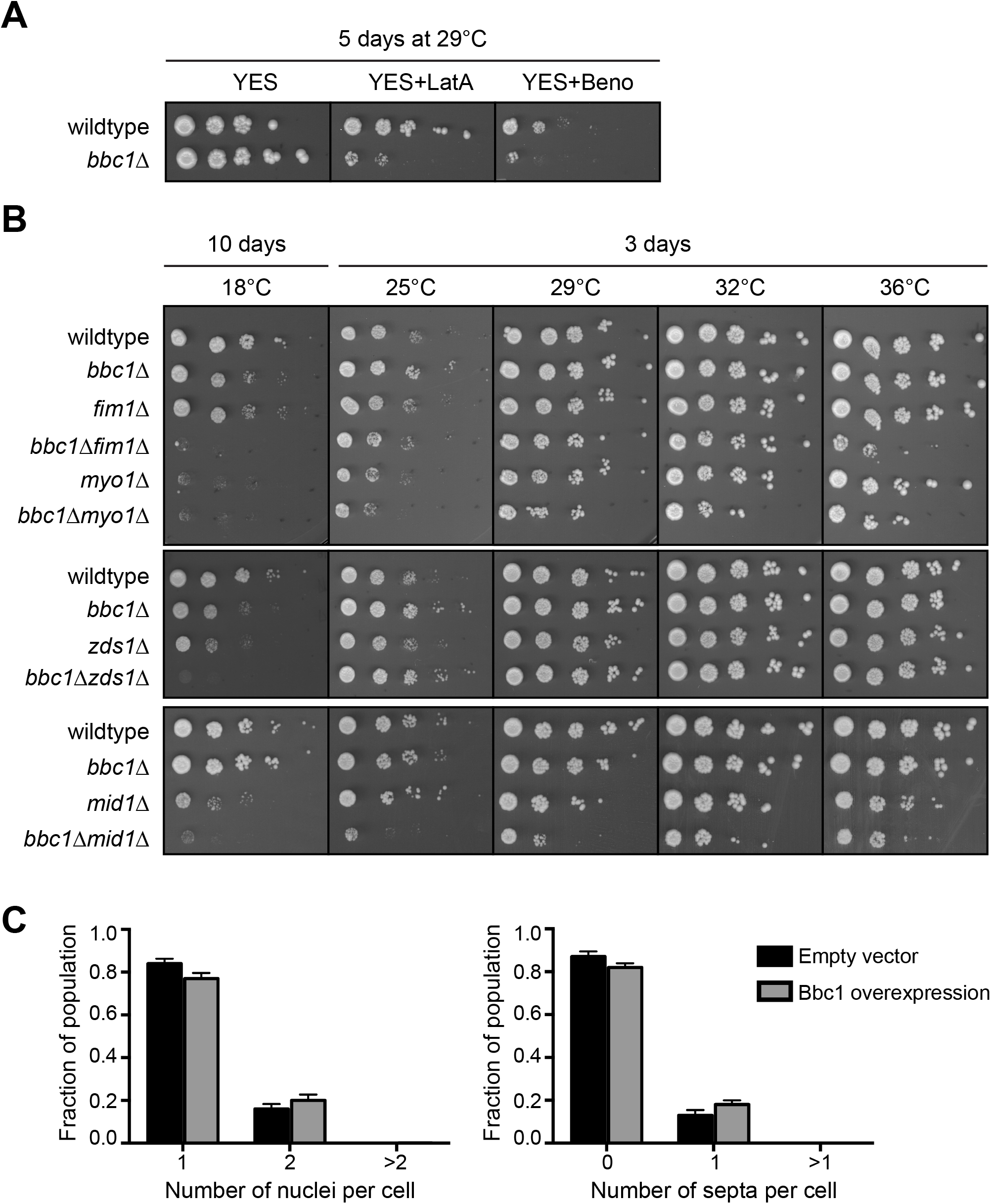
Drug sensitivities and genetic interactions of *bbc1*Δ suggest Bbc1 involvement in endocytosis. (A) Growth assays for drug sensitivity of WT and *bbc1Δ* cells. Cells were serially diluted, spotted onto YES plates or YES plates containing 0.25 μM latrunculin A or 10 μg/ml benomyl, and incubated for 5 days at 29°C. (B) Growth assays for genetic interactions of *bbc1Δ* with *fim1Δ, myo1Δ, zds1Δ*, and *mid1Δ*. Cells of indicated genotypes were serially diluted, spotted onto YES plates, and incubated for 3 days at indicated temperatures. (C) Quantification of number of nuclei (left) and number of septa (right) in WT cells transformed with the empty vector or Bbc1 overexpression construct (pREP3X-Bbc1-CDS). Cells were grown in EMM plus adenine and uracil in the absence of thiamine for 21-22 hours at 32°C to induce overexpression, fixed, and stained with DAPI and methyl blue. Graph represents average from 3 replicates, n ≥ 169 cells per replicate.

**FIGURE S2:**
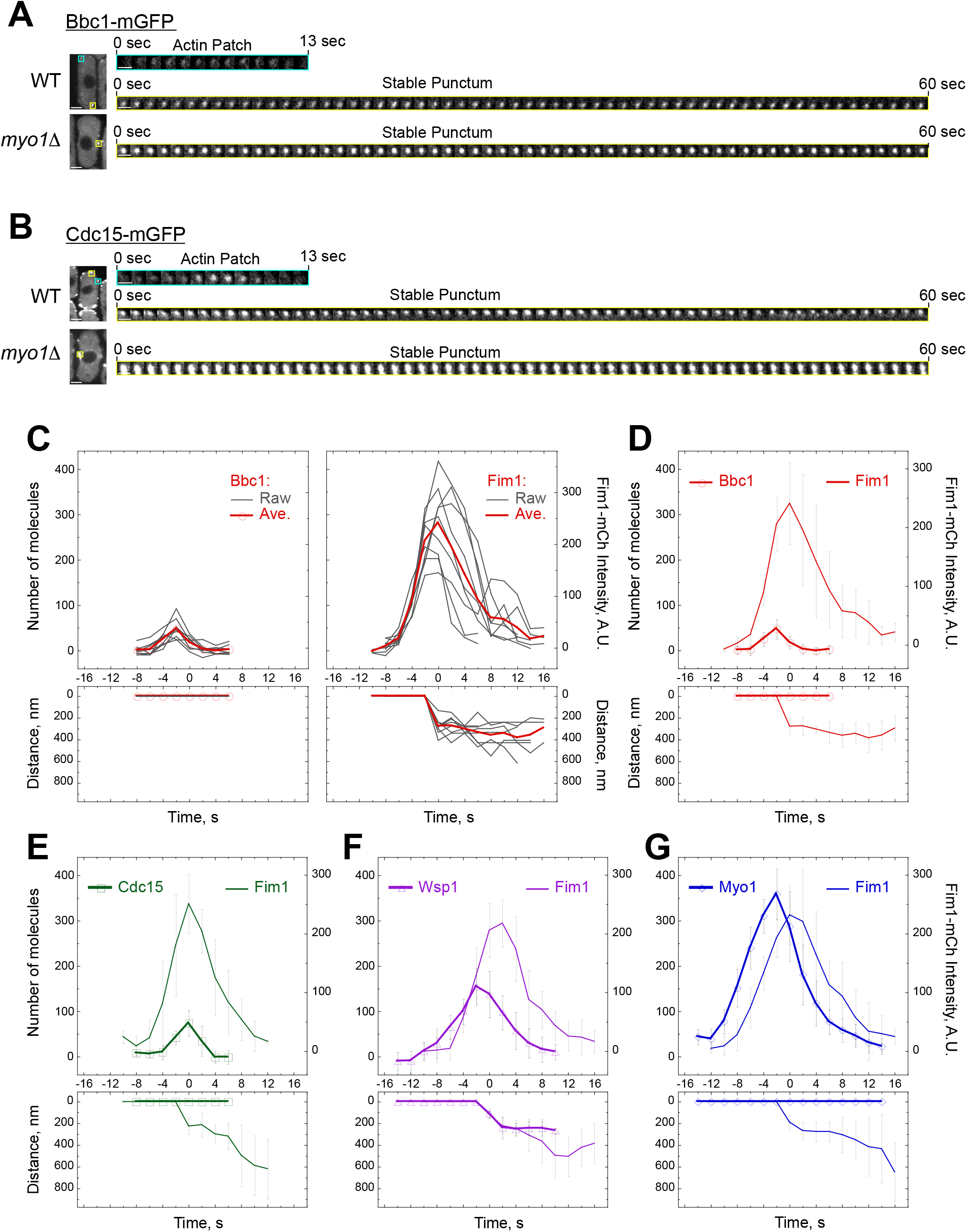
Bbc1 and Cdc15 localize to dynamic patches and stable puncta during interphase. (A-B) Confocal sections through the middle of single cells and time lapse montages of (A) Bbc1-mGFP and (B) Cdc15-mGFP in dynamic patches (boxed in teal) or stable puncta (boxed in yellow) in WT or *myo1*Δ cells. Images were taken at 1-second intervals. Scale bars, whole cells, 2 μm, montages, 1 μm. (C-G) Time courses of the number of molecules of mGFP-tagged Bbc1, Cdc15, Wsp1, and Myo1, Fim1-mCherry fluorescence intensity, and distance traveled by each marker in endocytic actin patches. These are the same as in Figure 2 but shown individually for each marker. (C) Raw (thin gray lines) and average time courses for Fim1-mCherry (red, no symbols) and Bbc1-mGFP (red circles). (D-G) Average time courses for (D) Bbc1-mGFP (red circles) and Fim1-mCherry (red, no symbols), (E) Cdc15-mGFP (green squares) and Fim1-mCherry (green, no symbols), (F) mGFP-Wsp1 (purple triangles) and Fim1-mCherry (purple, no symbols), and (G) mGFP-Myo1 (blue diamonds) and Fim1-mCherry (blue, no symbols). Cells expressing Fim1-mCherry with Bbc1-mGFP, Cdc15-mGFP, mGFP-Wsp1, or mGFP-Myo1 were imaged by spinning disk confocal microscopy in a medial plane at 2-second intervals. The intensity and position of each protein in patches were tracked and time courses for individual patches were aligned to the start of Fim1-mCherry movement (t=0). The intensity and distance values were averaged at each time point and mGFP intensity values were converted into the number of molecules based on previously measured numbers of molecules (Sirotkin et al., 2010). Error bars represent SD.

**FIGURE S3:**
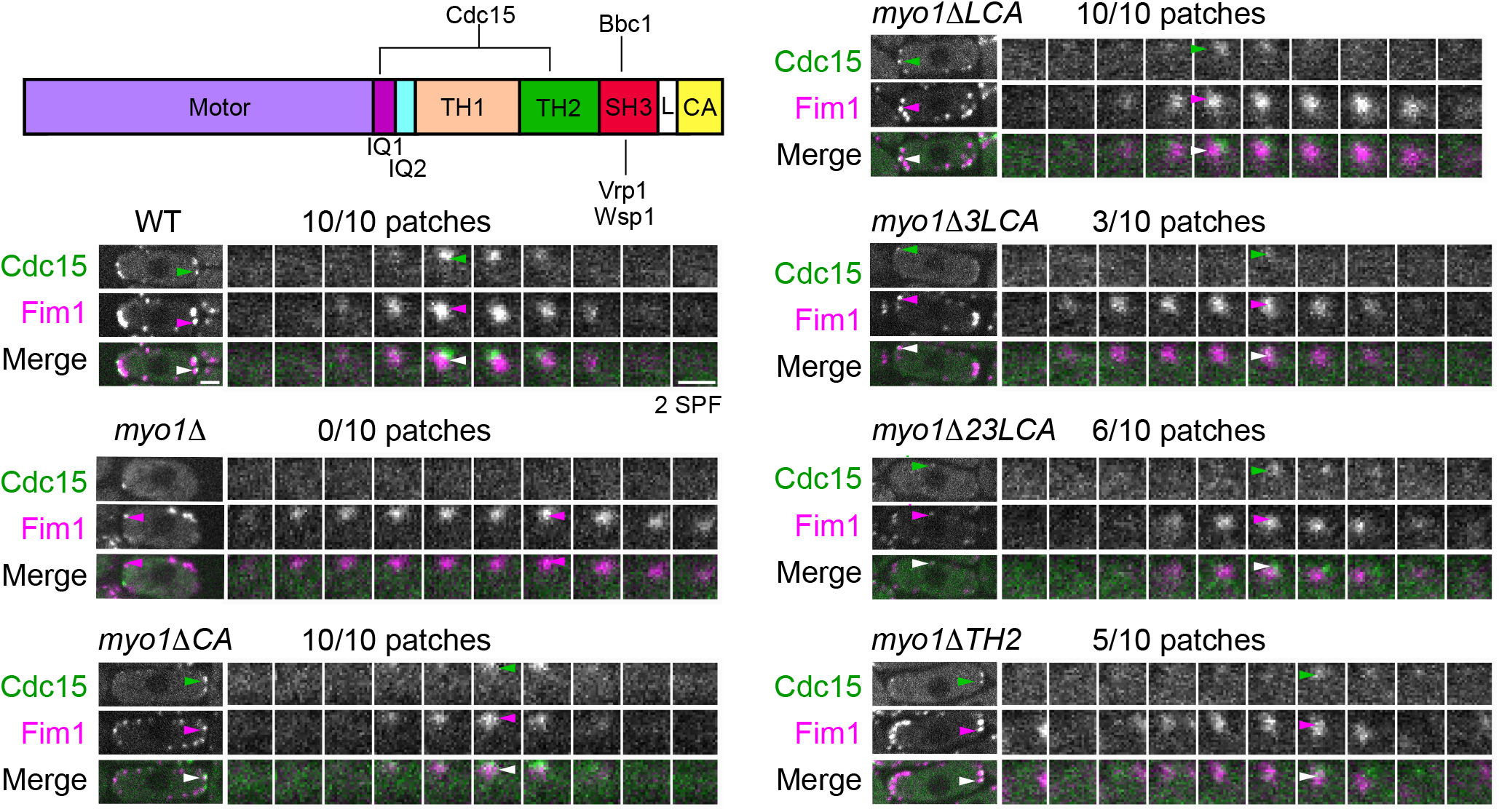
The Myo1 TH2 and SH3 domains contribute to Cdc15 patch recruitment. A schematic diagram of the Myo1 protein domains with relevant binding partners and example montages of co-localization analysis of Cdc15-mGFP (green) and Fim1-mCherry (magenta) in patches in WT and indicated Myo1 tail domain deletion strains. Montages show the protein localization dynamics at intervals of 2 SPF in patches indicated by arrowheads on montages and whole cell images on the left. Numbers show the fraction of Fim1 patches that also contained Cdc15. Scale bars, whole cells, 2 μm; montages, 1 μm.

**FIGURE S4:**
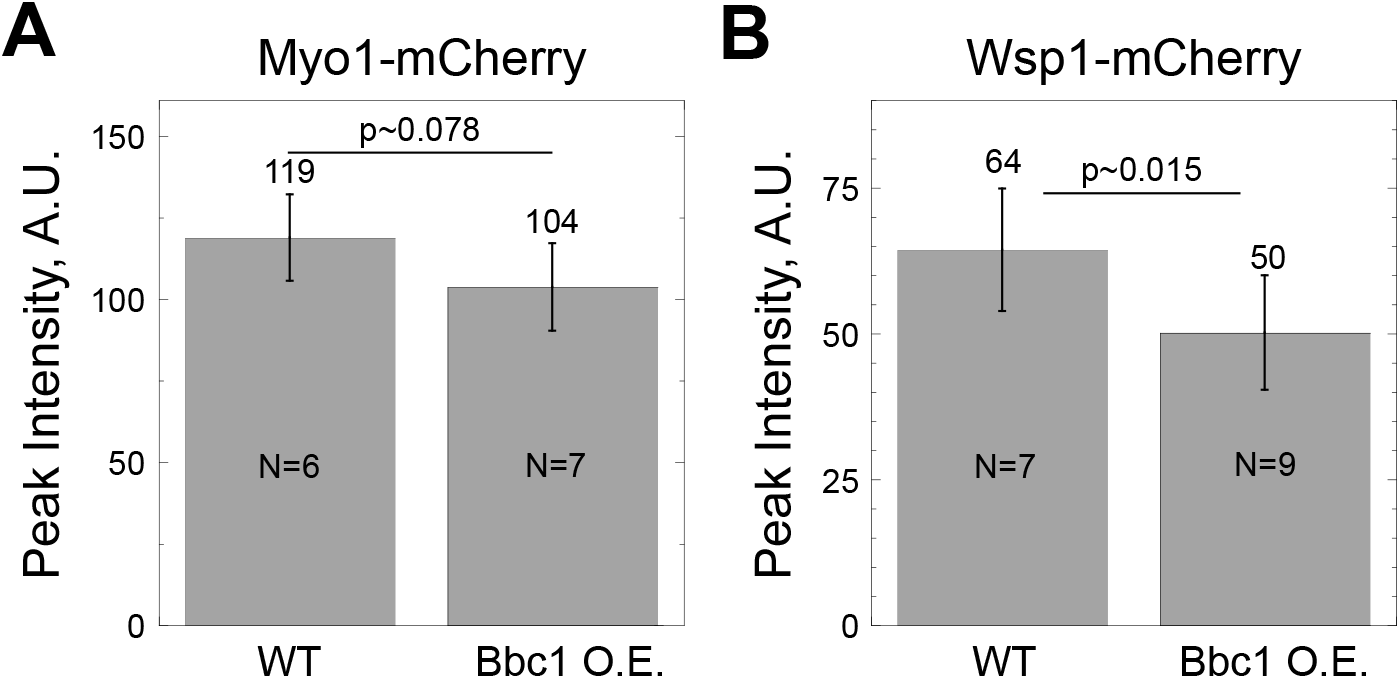
Bbc1 overexpression reduces Myo1 and Wsp1 accumulation in patches. (A-B) Bar graphs represent the average peak intensities of (A) Myo1-mCherry and (B) Wsp1-mCherry in patches in WT and mGFP-Bbc1-overexpressing cells. Error bars represent SD. N indicates number of patches analyzed. P values represent statistical significance as determined by t-test.

**FIGURE S5:**
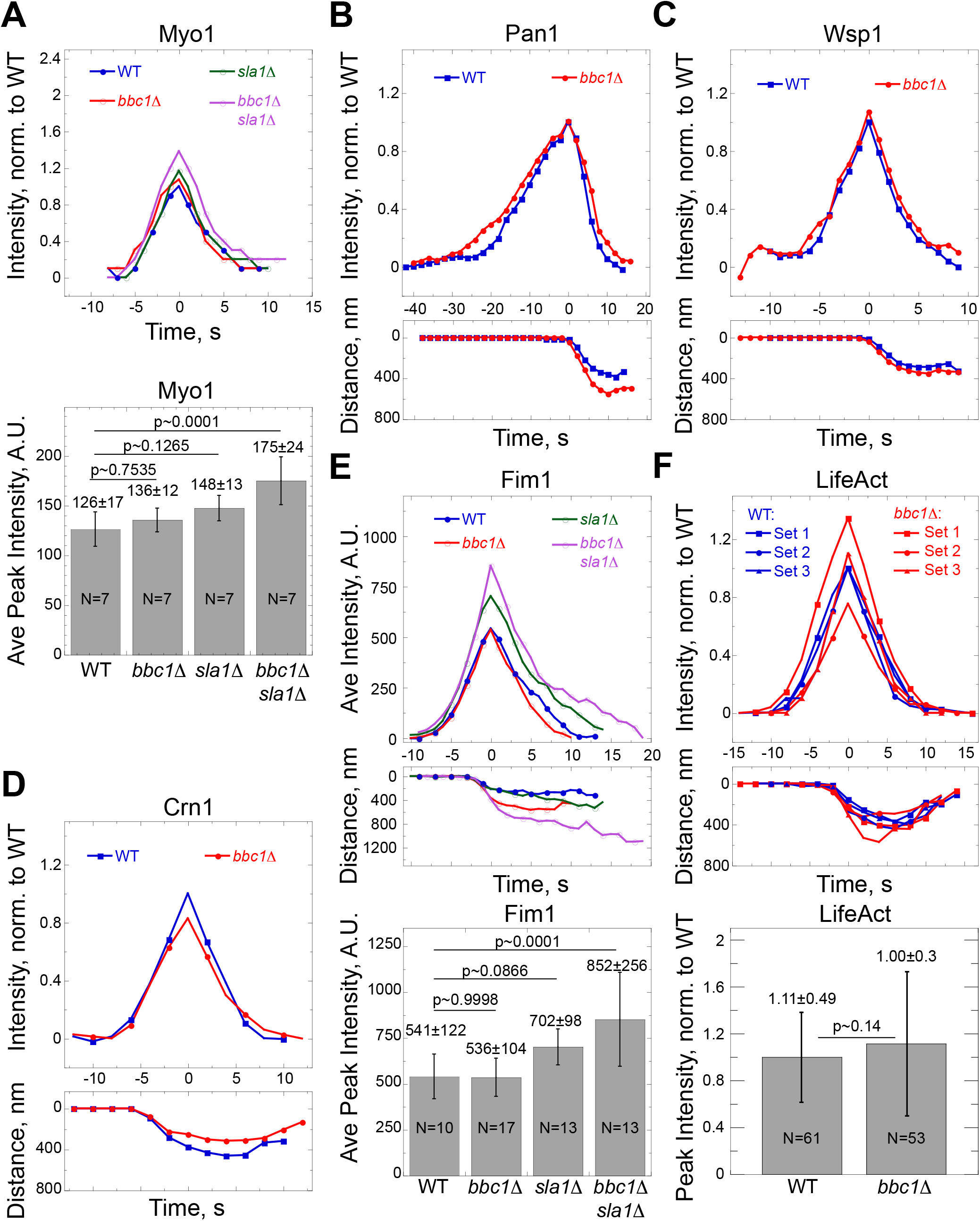
Effects of *bbc1Δ* and *sla1Δ* on patch accumulation of endocytic proteins. (A-F) Average time courses of intensities, (B-F) distances traveled, and (A, E, F) average peak intensities for (A) mGFP-Myo1, (B) Pan1-GFP, (C) mGFP-Wsp1, (D) Crn1-GFP, (E) Fim1-mGFP, and (F) LifeAct-mCherry in endocytic patches in WT (blue), *bbc1Δ* (red), *sla1Δ* (green), and *bbc1Δ sla1Δ* (purple) cells. Time courses for individual patches were aligned to the time each patch reached peak intensity (t=0), averaged at each time point, and, except for (E), normalized to peak intensity in WT cells. N= 7 patches per condition in (A) and N= at least 10 patches per condition in (B-E). In (F), LifeAct-mCherry patch dynamics were measured in WT (blue) or *bbc1Δ* (red) cells also expressing Myo1-GFP (Set 1), Crn1-GFP (Set 2), or Pan1-GFP (Set 3) and normalized peak intensities for individual patches from all datasets were combined and averaged for a total of N=61 patches in WT and N=53 patches in *bbc1Δ* cells. Error bars on bar graphs represent SD. P values represent statistical significance as determined by (A, E) ANOVA with Tukey’s Post-Hoc test or (F) t-test.

**FIGURE S6.**
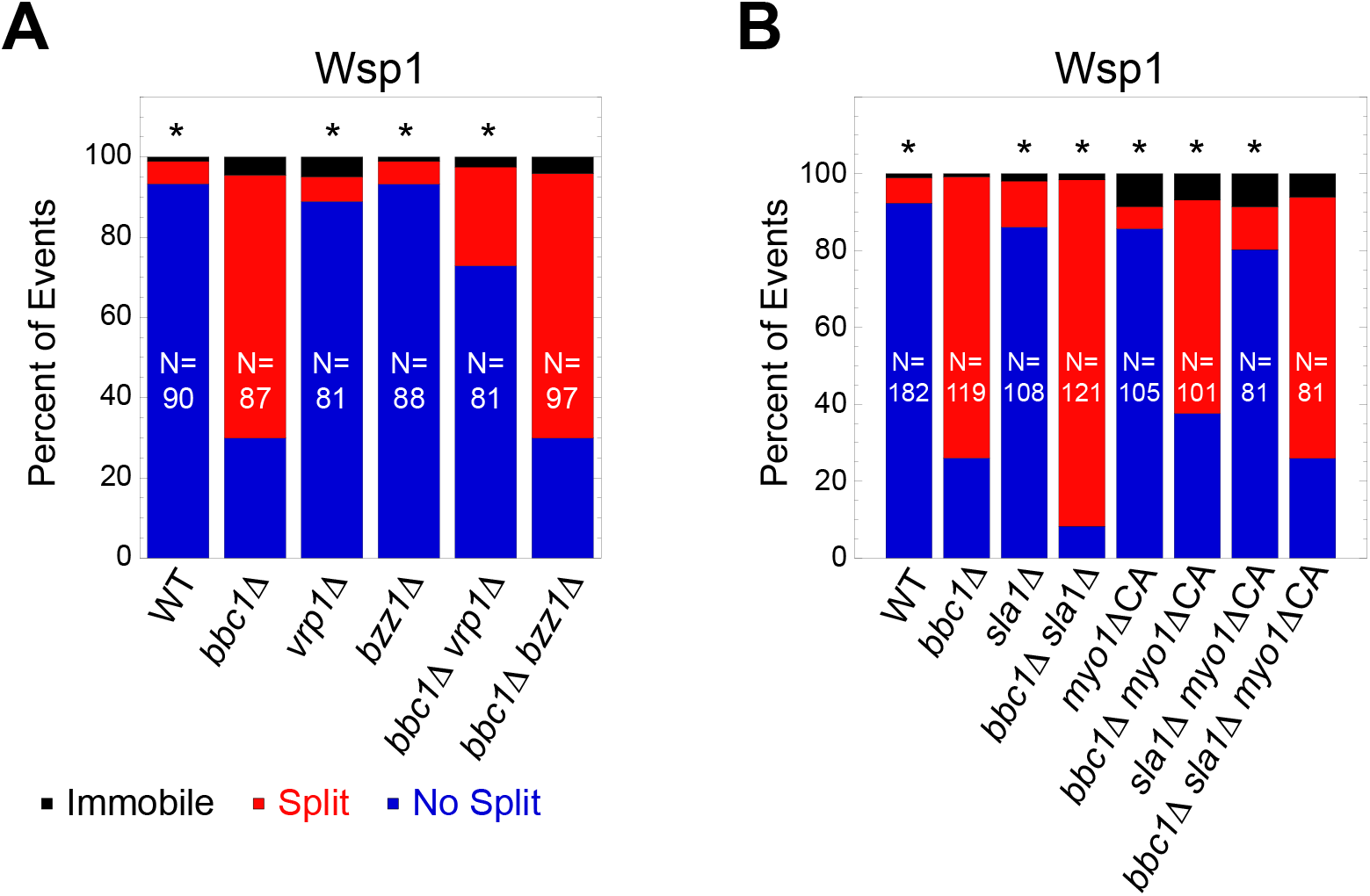
The *bbc1*Δ alone is sufficient for Vrp1- and Myo1 SH3 domain-mediated Wsp1 patch splitting. Wsp1 patch splitting and its genetic dependencies are similar in *bbc1*Δ and *bbc1*Δ *sla1*Δ cells (compare to Figure 6). (A-B) Quantification of the percent of events when mGPF-Wsp1 patches internalized normally (blue), remained immobile (black), or split in two (red) in indicated strains with different combinations of (A) *bbc1Δ* and *vrp1Δ* or *bzz1Δ* and (B) *bbc1Δ, sla1Δ*, and *myo1ΔCA*. Asterisks indicate a significant difference from *bbc1Δ* as determined by Chi-squared test, *p<0.05. N indicates number of patches analyzed.

**FIGURE S7:**
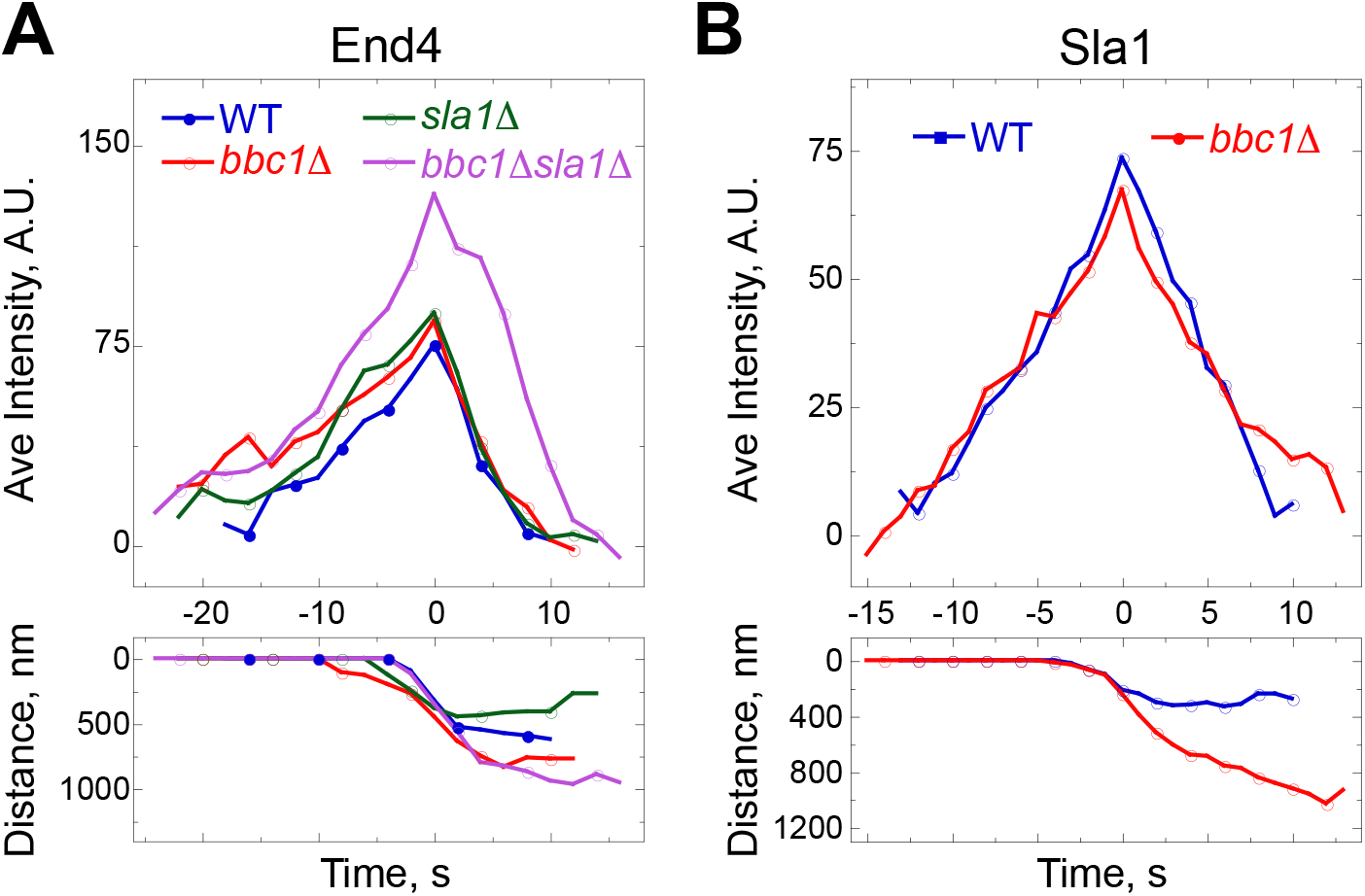
The *bbc1Δ* increases the distance traveled but not accumulation of End4 and Sla1 in patches. (A-B) Average time courses of intensities and distances traveled for (A) End4-mGFP in endocytic patches in WT (blue), *bbc1Δ* (red), *sla1Δ* (green), and *bbc1Δ sla1Δ* (purple) cells, and (B) Sla1-mGFP in patches in WT (blue) and *bbc1Δ* (red) cells. Time courses for individual patches were aligned to the time each patch reached peak intensity (t=0) and averaged at each time point. (A) N=5 and (B) N= 7-10 patches per condition.

### SUPPLEMENTAL VIDEO LEGENDS

Video 1: Myo1 associates with the tip of actin plumes in *bbc1 Δ sla1 Δ* cells. Time series of images of Myo1-GFP (left, green in merge), LifeAct-mCherry (middle, magenta in merge), and the merge (right) in a single confocal section through the middle of live *bbc1 Δ sla1 Δ* cells. Images were acquired at 2-second intervals for 60 seconds and are displayed at the rate of 7 frames per second.

Video 2: Myo1 association with actin plumes is dynamic and long-lived. Time series of images of mGFP-Myo1 in a single confocal section through the middle of live *sla1 Δ* (left) and *bbc1 Δ sla1 Δ* (right) cells. Images were acquired at 1 frame per second and are displayed at the rate of 7 frames per second.

Video 3: Wsp1 patches split in the absence of Bbc1. Time series of images of mGFP-Wsp1 in in a single confocal section through the middle of live (from left to right) WT, *bbc1 Δ, sla1 Δ*, and *bbc1 Δ sla1 Δ* cells. Images were acquired at 1 frame per second and are displayed at the rate of 7 frames per second.

### SUPPLEMENTAL TABLES

**Table S1.** Proteins associated with Bbc1. LC-MS/MS results with the number of unique peptides and percent sequence coverage provided.

| Purification | Protein | Unique peptides | % Coverage |
| --- | --- | --- | --- |
| Cyk3-TAP | Cyk3 | 27 | 39 |
|  | Cdc15 | 40 | 47 |
|  | Bbc1 | 4 | 4 |
| Fic1-GFP | Fic1 | 34 | 56 |
|  | Cdc15 | 31 | 40 |
|  | Bbc1 | 13 | 13 |
| Bbc1-TAP | Bbc1 | 187 | 61 |
|  | Myo1 | 60 | 55 |
|  | Pan1 | 44 | 30 |
|  | Cdc15 | 26 | 34 |

Abbreviations: TAP, tandem affinity purification; GFP, green fluorescent protein.

**Table S2.**
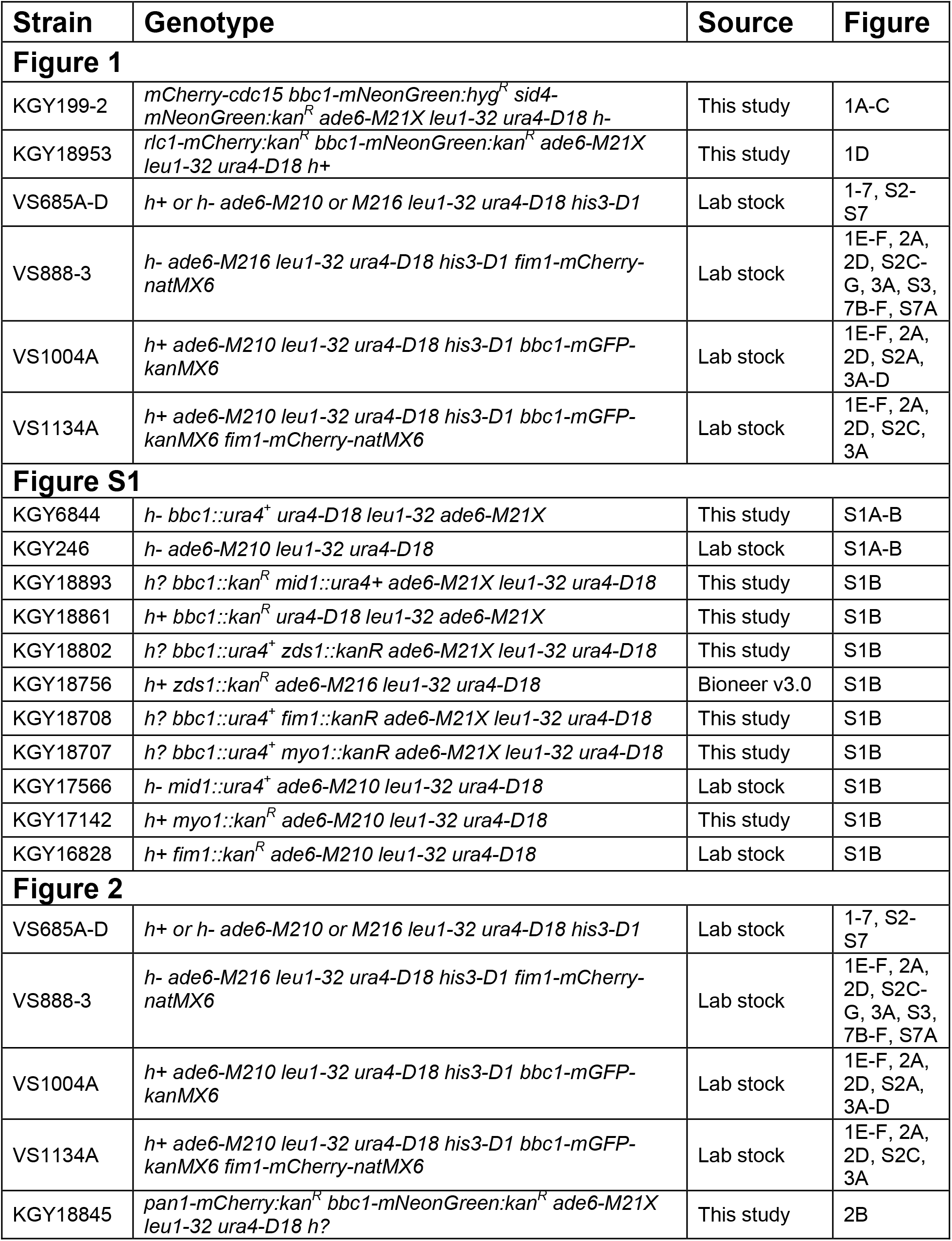
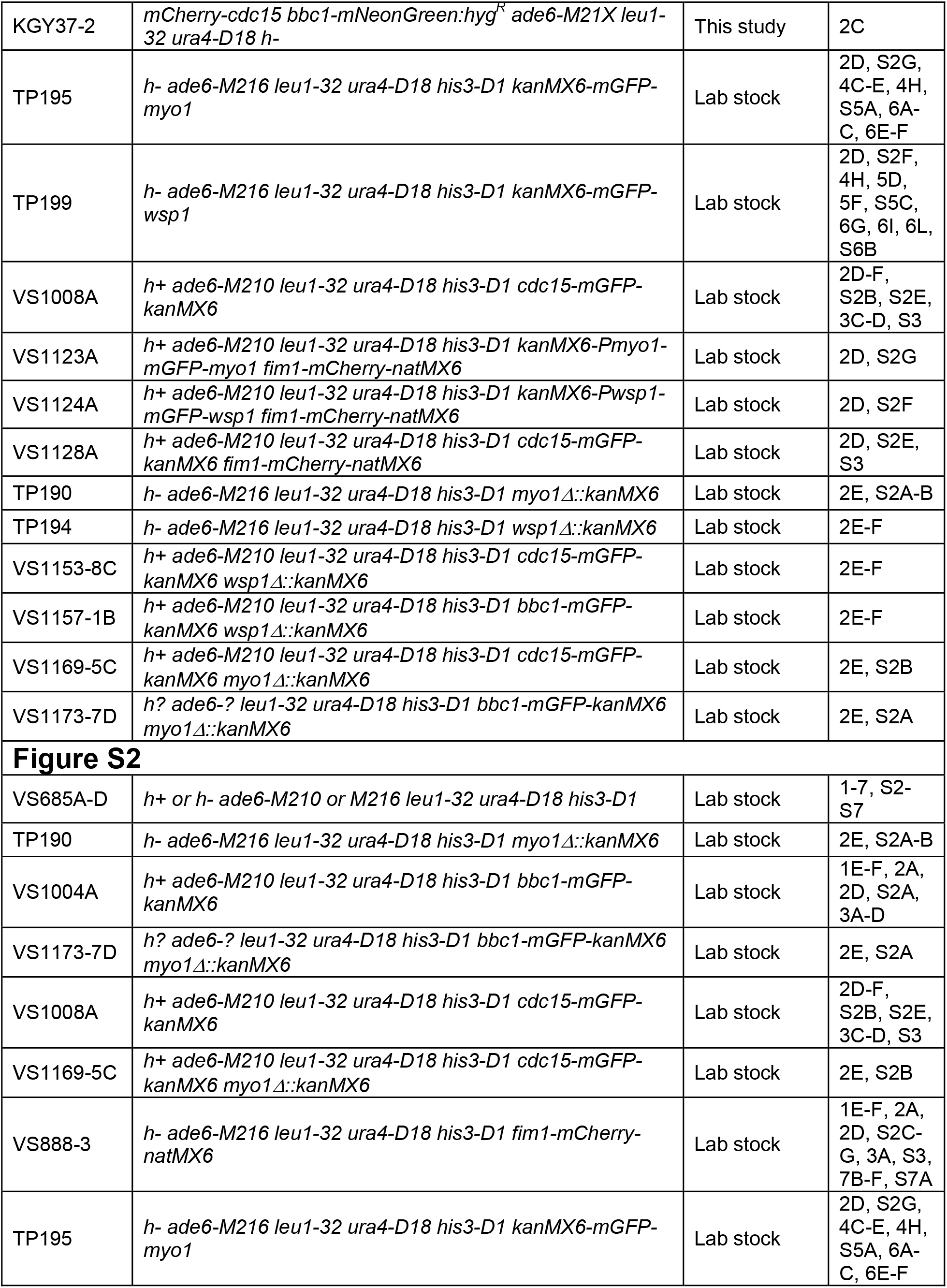
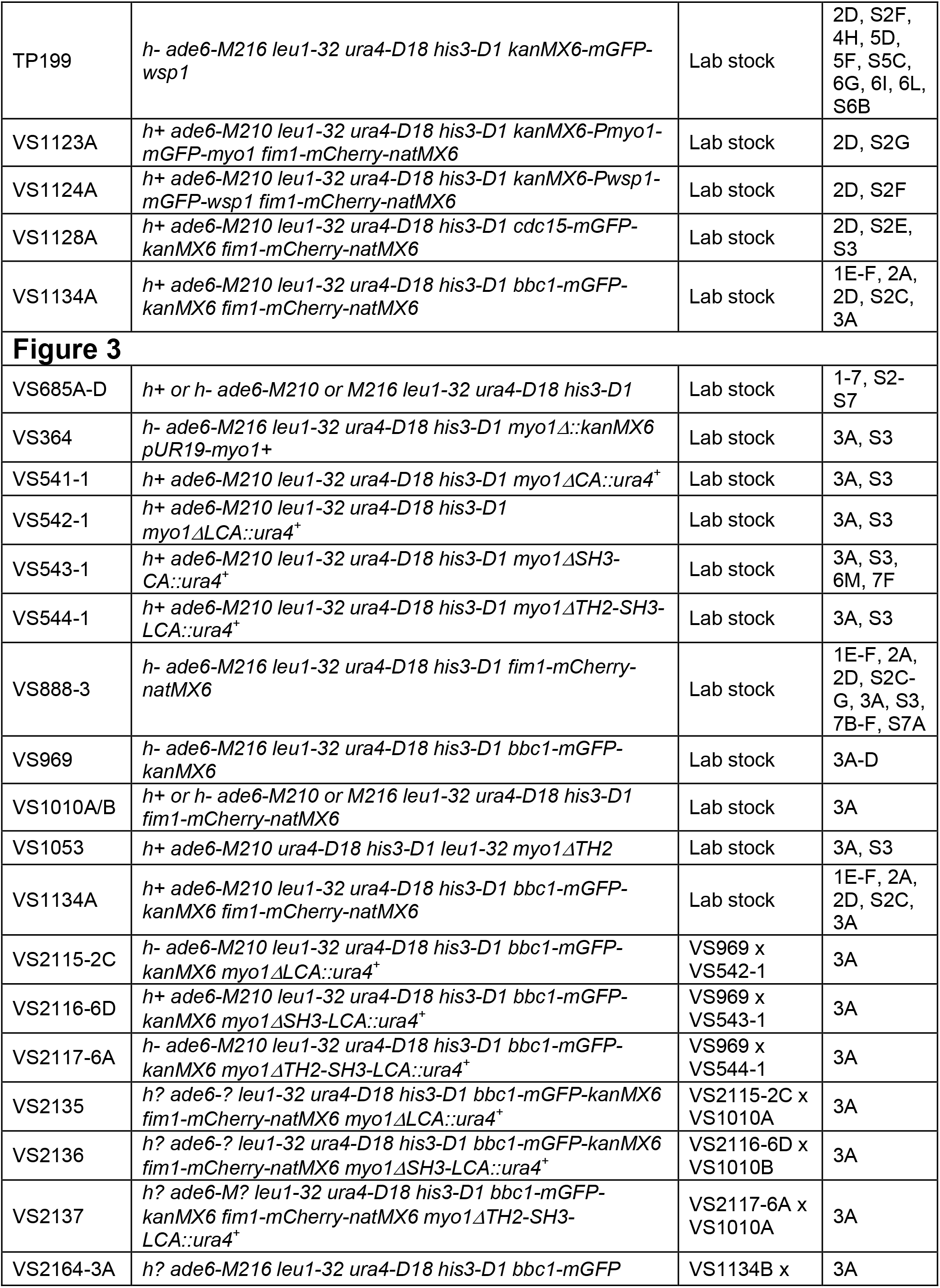
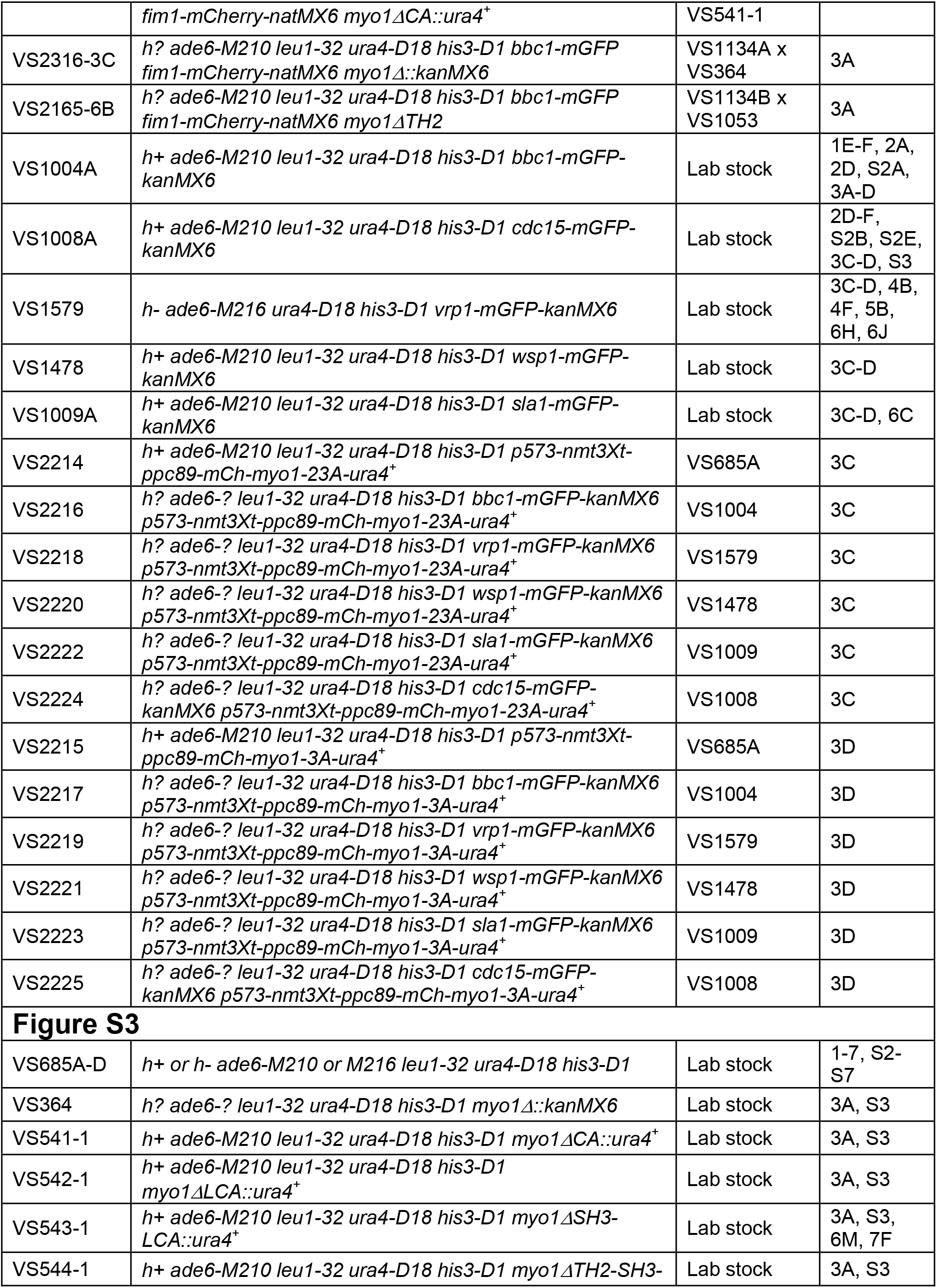
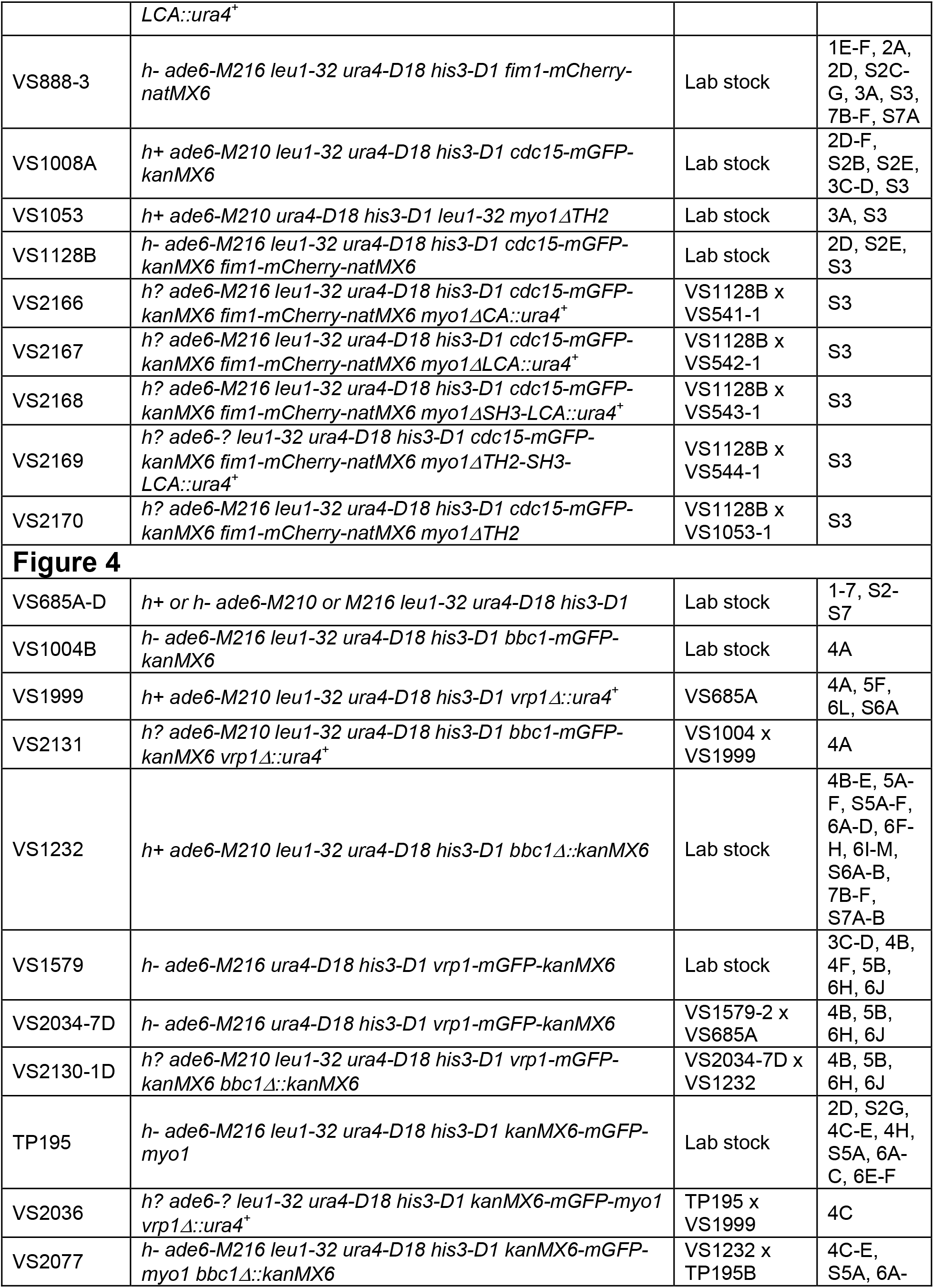
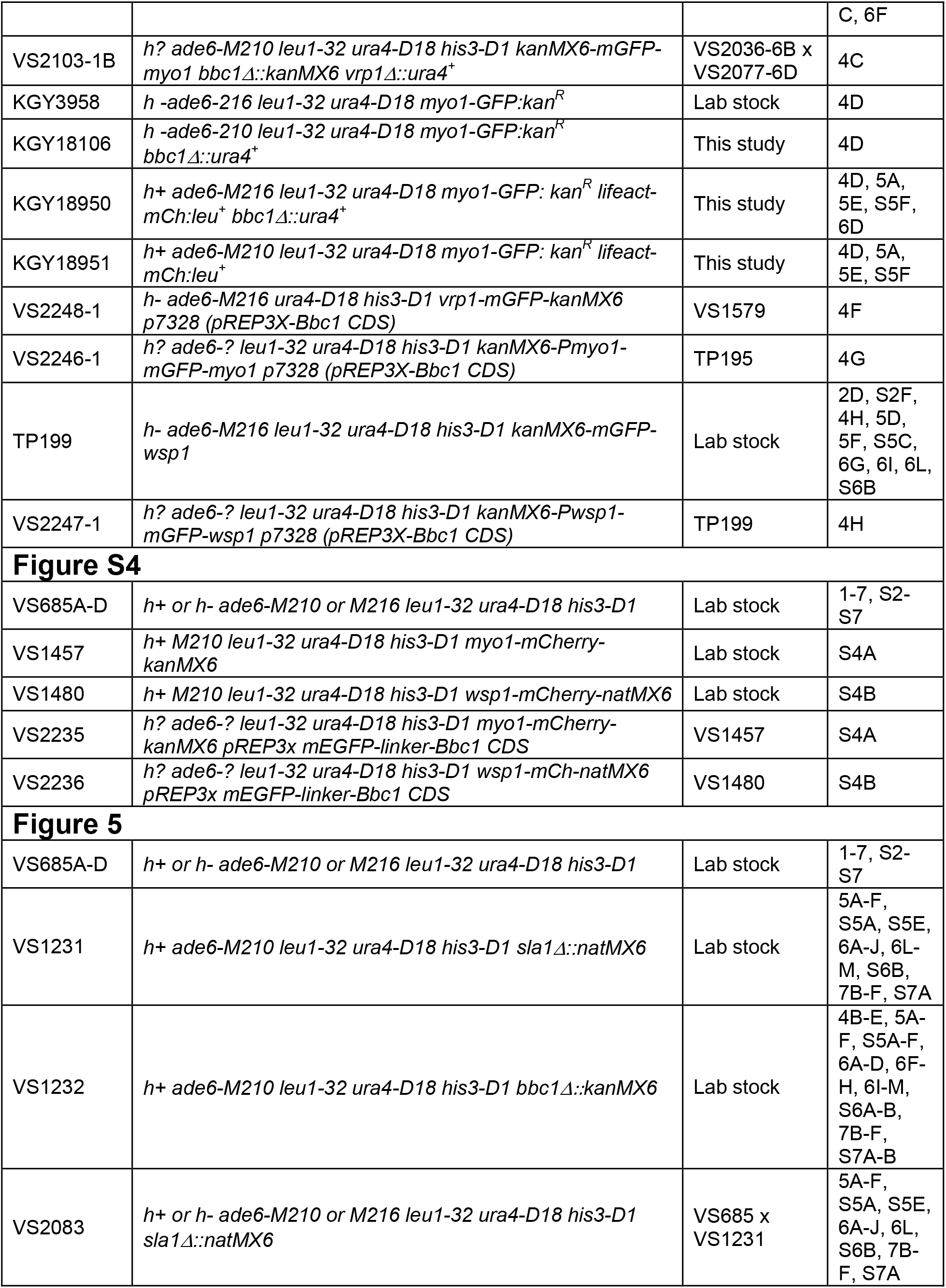
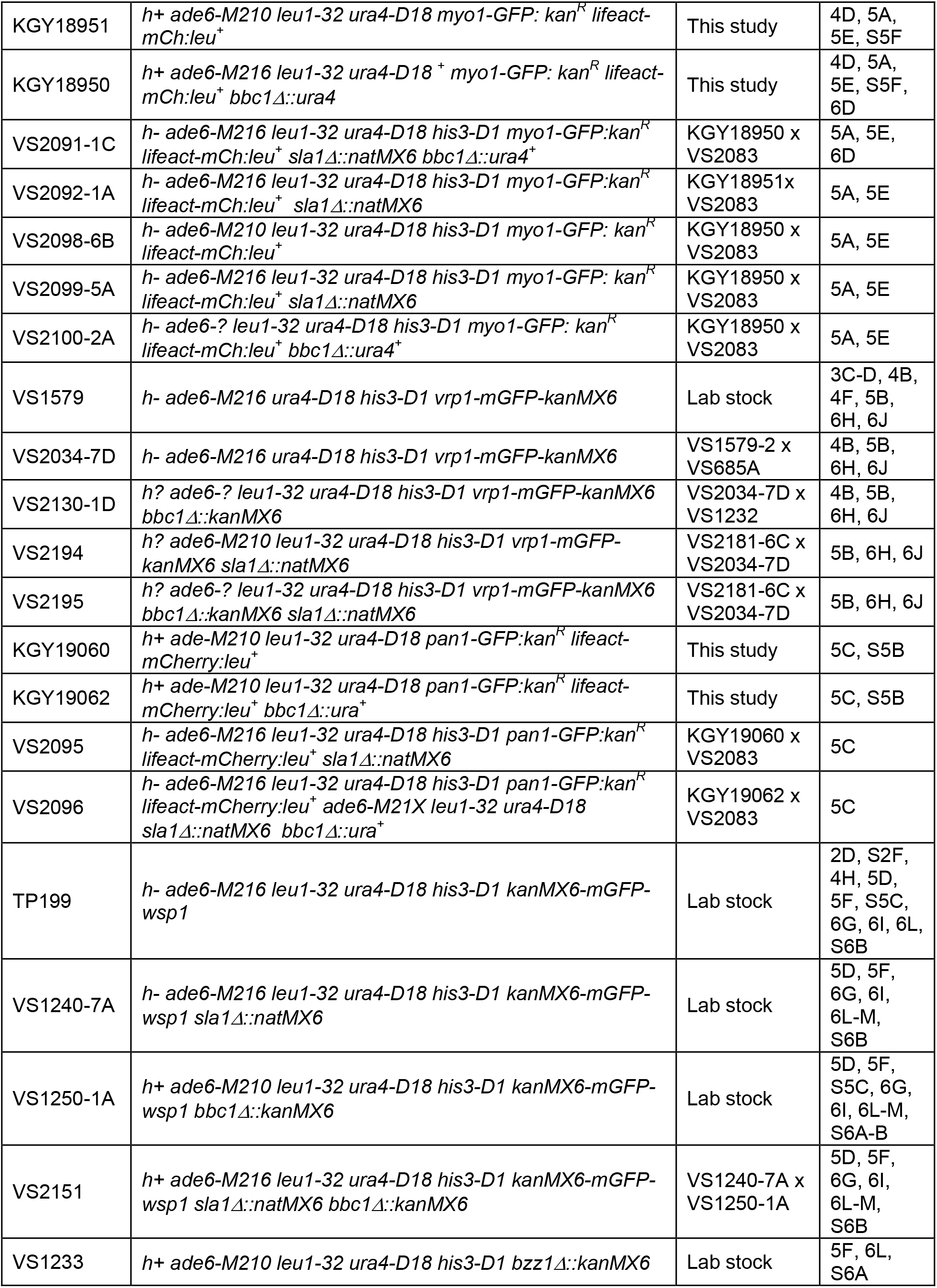
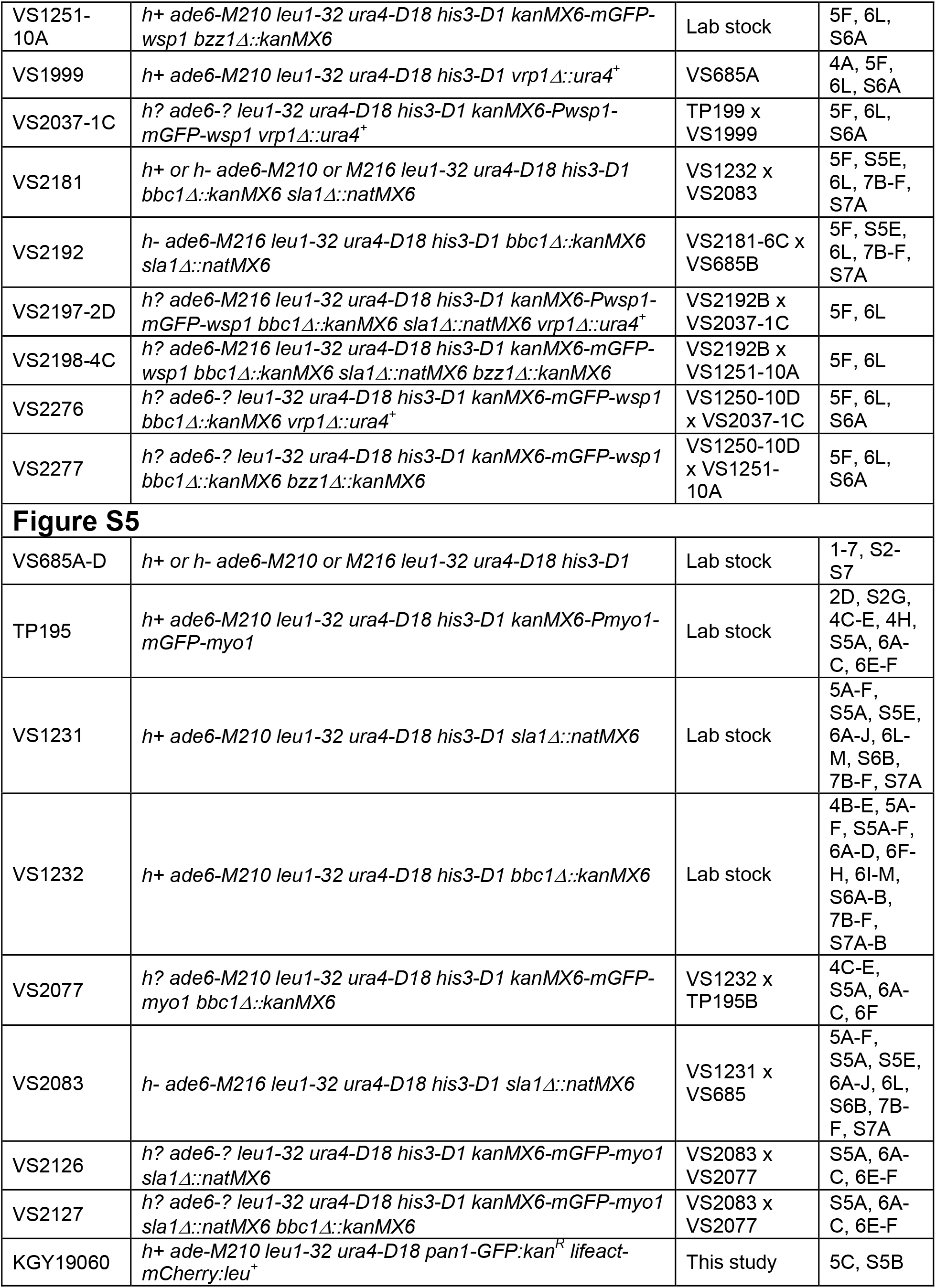
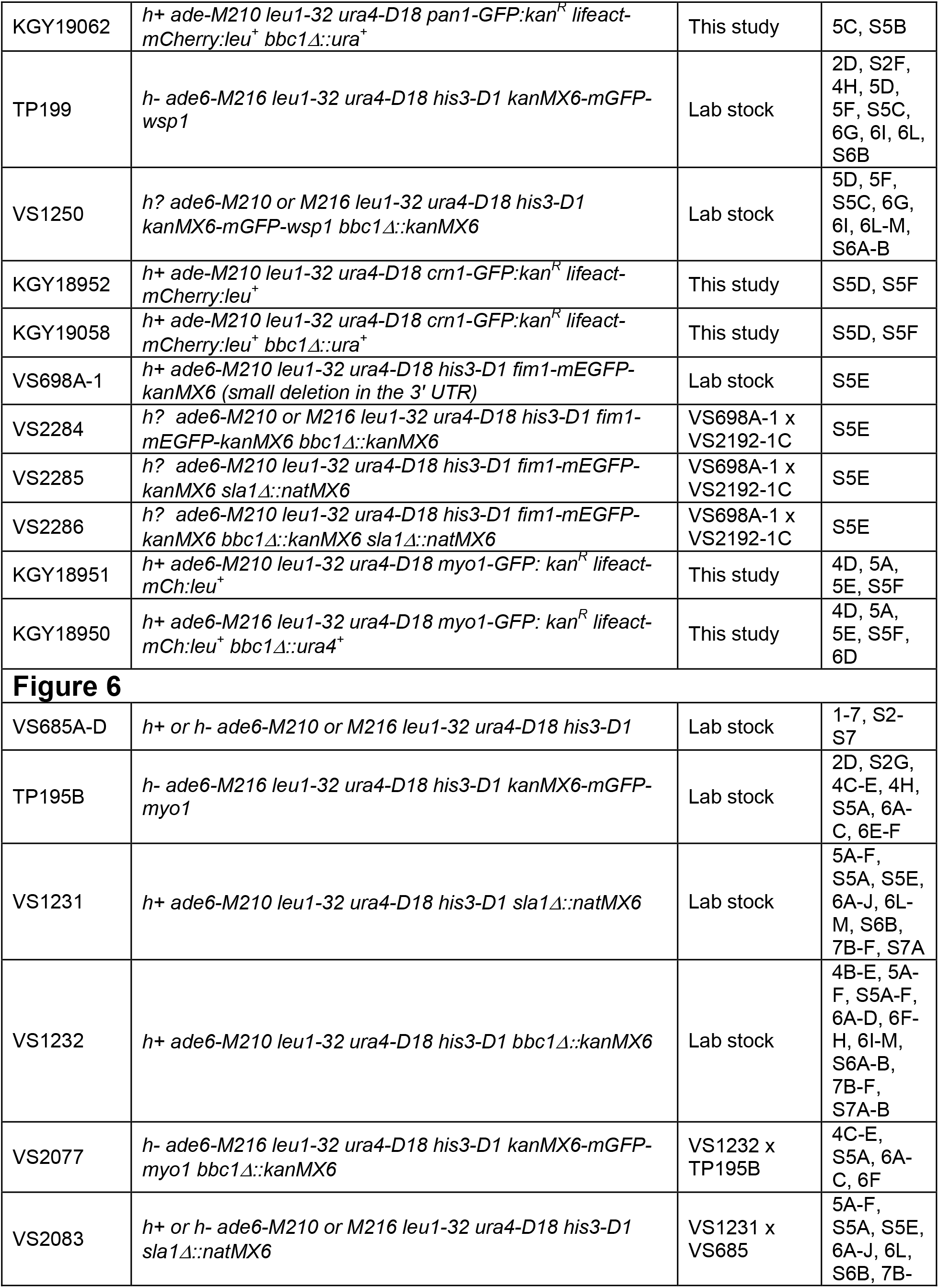
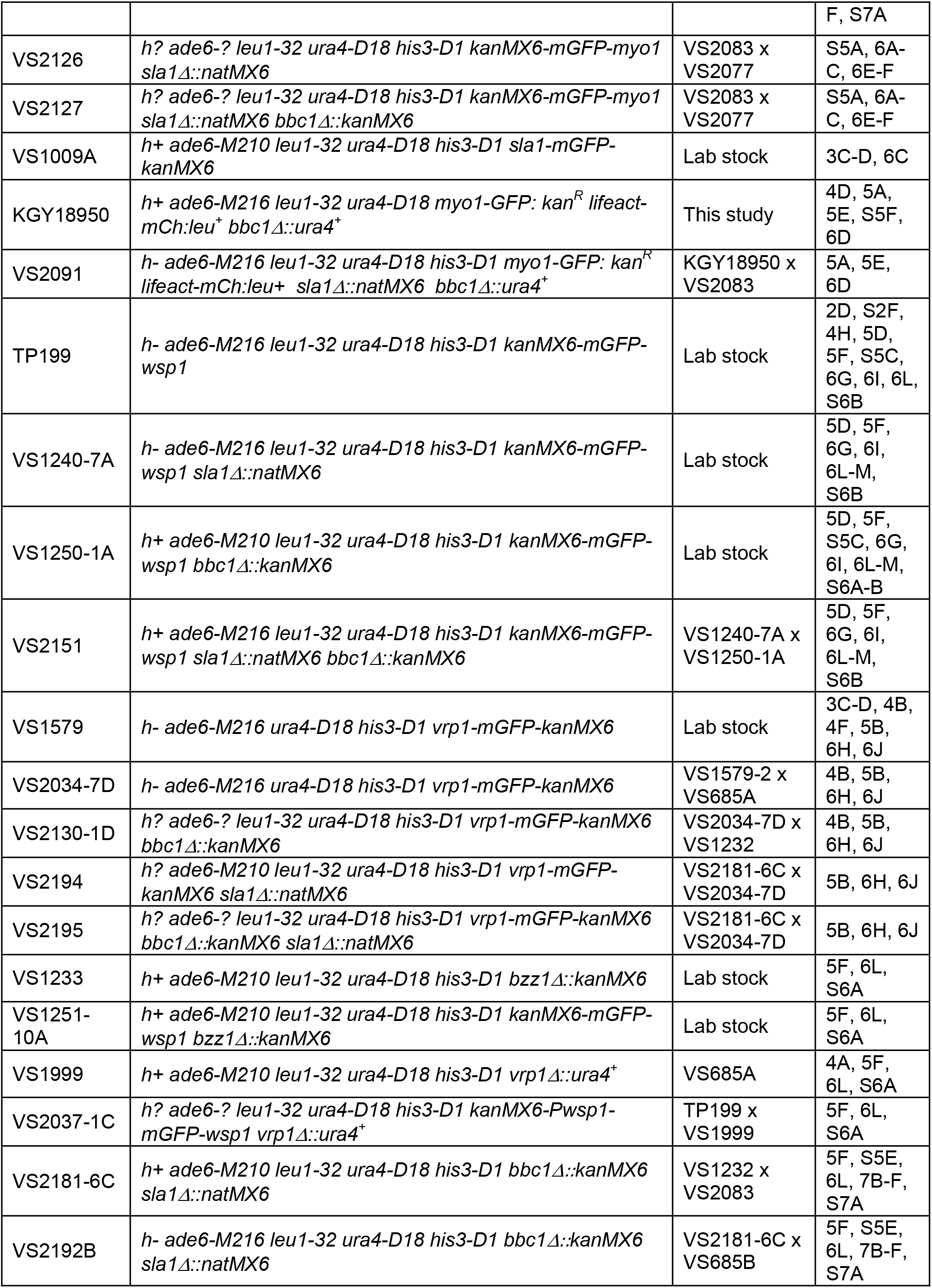
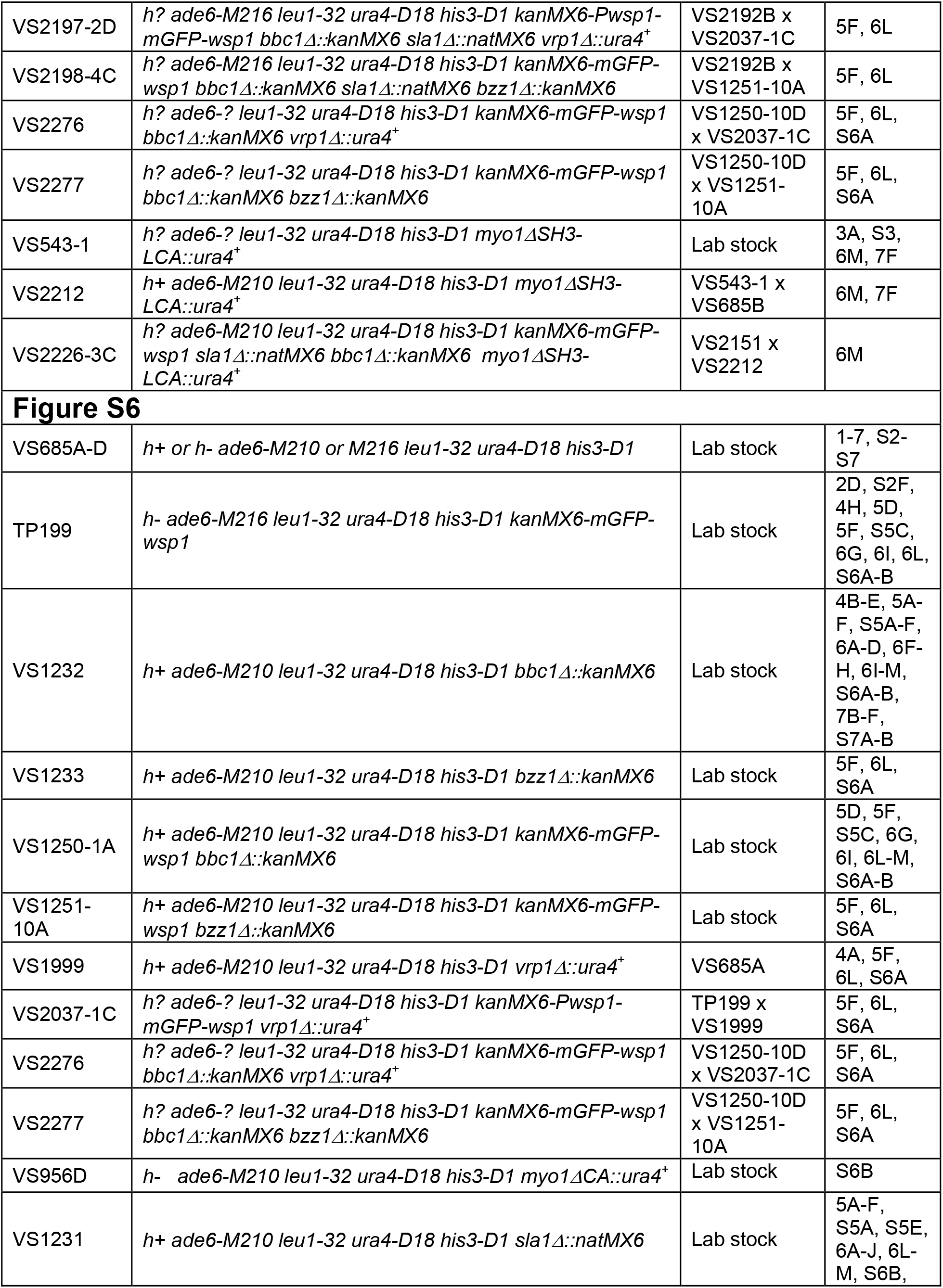
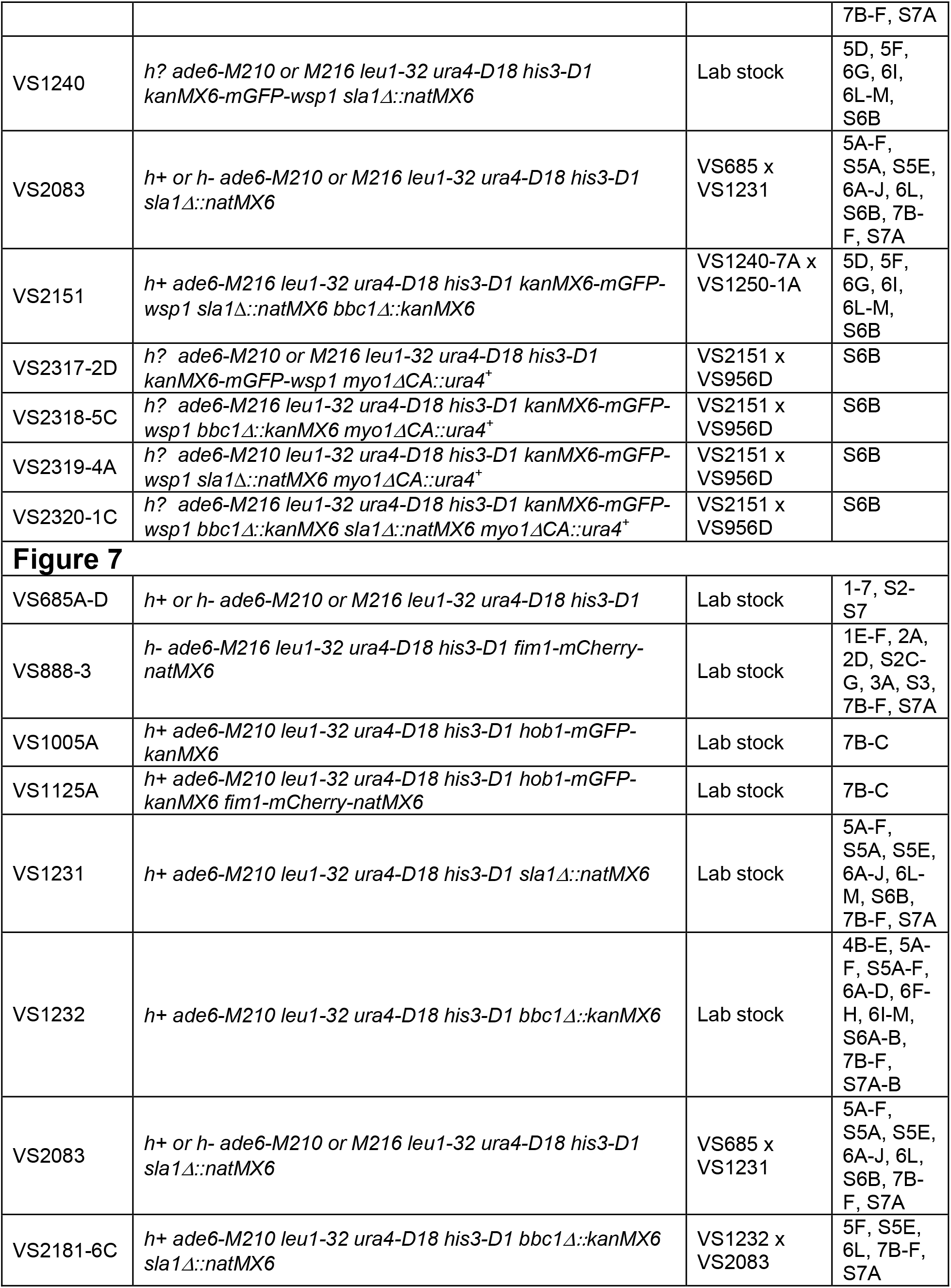
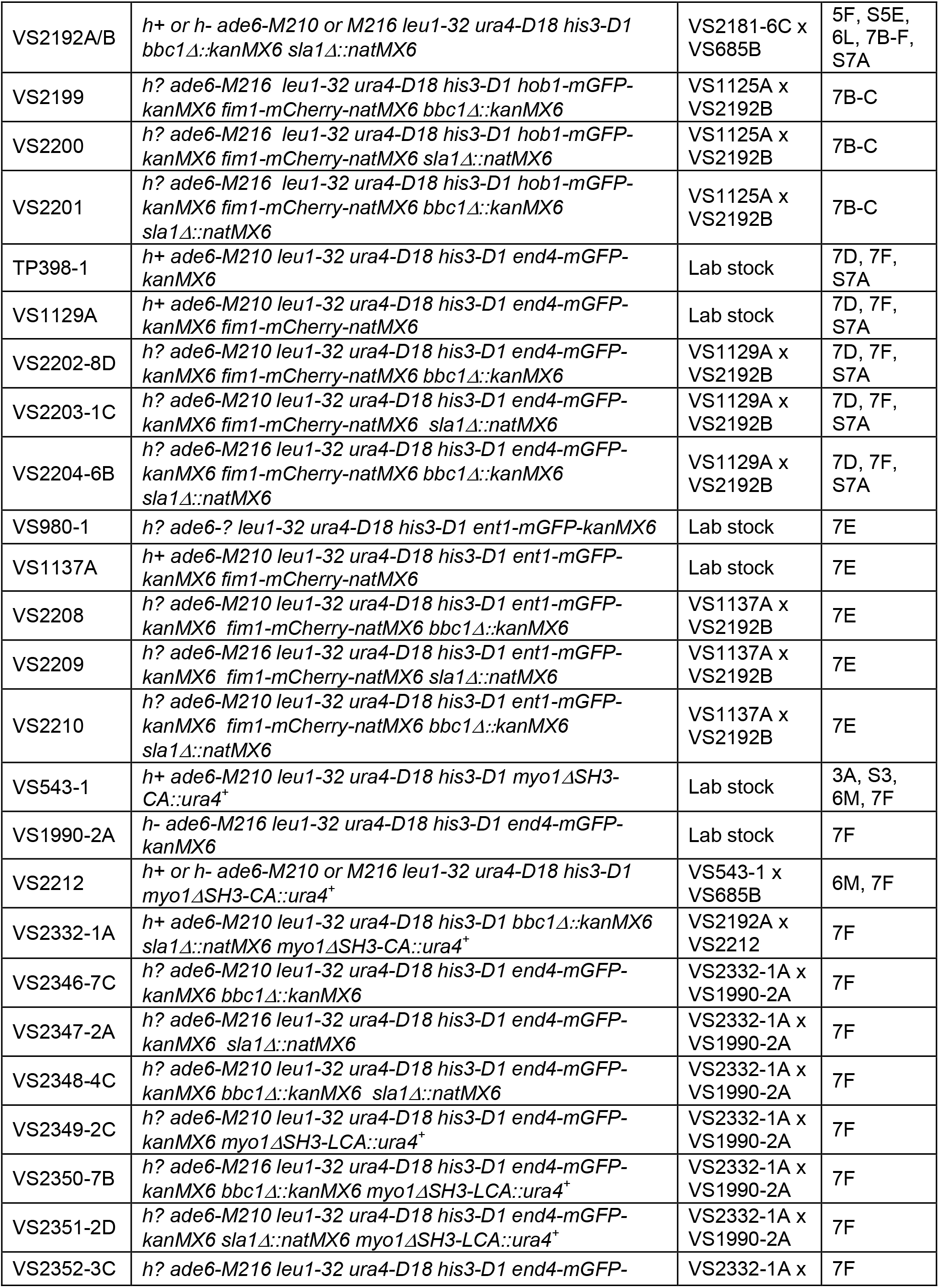
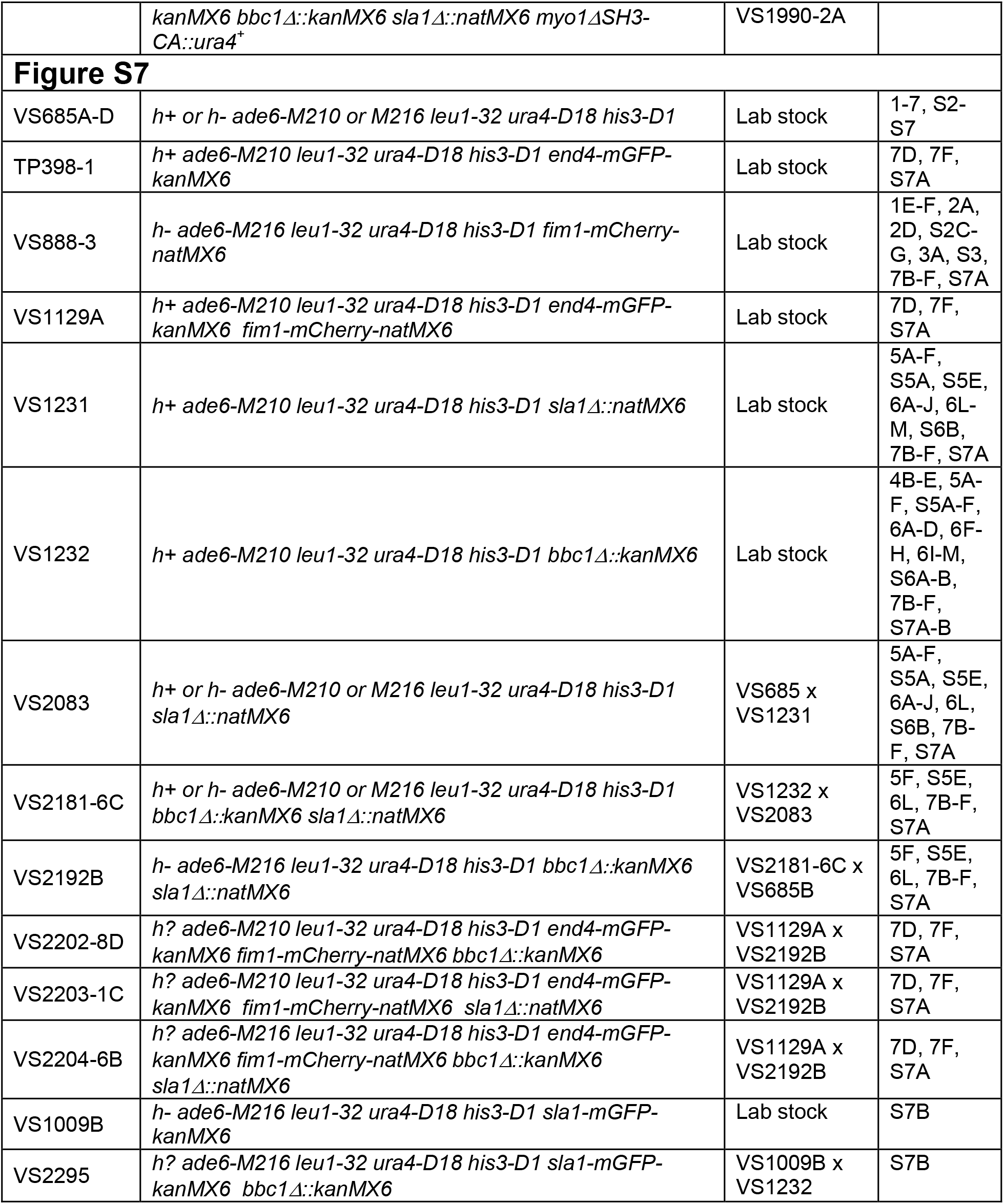
*S. pombe* strains used in this study.

## REFERENCES

Aghamohammadzadeh, S., and K.R. Ayscough. 2009. Differential requirements for actin during yeast and mammalian endocytosis. Nat Cell Biol. 11:1039–1042.

Anderson, B.L., I. Boldogh, M. Evangelista, C. Boone, L.A. Greene, and L.A. Pon. 1998. The Src homology domain 3 (SH3) of a yeast type I myosin, Myo5p, binds to verprolin and is required for targeting to sites of actin polarization. J Cell Biol. 141:1357–1370.

Arasada, R., and T.D. Pollard. 2011. Distinct roles for F-BAR proteins Cdc15p and Bzz1p in actin polymerization at sites of endocytosis in fission yeast. Current Biology. 21:1450–1459.

Arasada, R., W.A. Sayyad, J. Berro, and T.D. Pollard. 2018. High-speed superresolution imaging of the proteins in fission yeast clathrin-mediated endocytic actin patches. Molecular Biology of the Cell. 29:235–376.

Bahler, J., J.Q. Wu, M.S. Longtine, N.G. Shah, A. McKenzie, 3rd, A.B. Steever, A. Wach, P. Philippsen, and J.R. Pringle. 1998. Heterologous modules for efficient and versatile PCR-based gene targeting in Schizosaccharomyces pombe. Yeast. 14:943–951.

Barker, S.L., L. Lee, B.D. Pierce, L. Maldonado-Báez, D.G. Drubin, and B. Wendland. 2007. Interaction of the Endocytic Scaffold Protein Pan1 with the Type I Myosins Contributes to the Late Stages of Endocytosis. Molecular Biology of the Cell. 18:2893–2903.

Basu, R., and F. Chang. 2011. Characterization of dip1p reveals a switch in Arp2/3-dependent actin assembly for fission yeast endocytosis. Curr Biol. 21:905–916.

Basu, R., E.L. Munteanu, and F. Chang. 2014. Role of turgor pressure in endocytosis in fission yeast. Molecular Biology of the Cell. 25:679–687.

Berro, J., V. Sirotkin, and T.D. Pollard. 2010. Mathematical modeling of endocytic actin patch kinetics in fission yeast: disassembly requires release of actin filament fragments. Molecular Biology of the Cell. 21:2905–2915.

Boettner, D.R., R.J. Chi, and S.K. Lemmon. 2012. Lessons from yeast for clathrin-mediated endocytosis. Nat Cell Biol. 14:2–10.

Boettner, D.R., H. Friesen, B. Andrews, and S.K. Lemmon. 2011. Clathrin light chain directs endocytosis by influencing the binding of the yeast Hip1R homologue, Sla2, to F-actin. Molecular Biology of the Cell. 22:3699–3714.

Boulant, S., C. Kural, J.-C. Zeeh, F. Ubelmann, and T. Kirchhausen. 2011. Actin dynamics counteract membrane tension during clathrin-mediated endocytosis. Nature Cell Biology. 13:1124–1131.

Carnahan, R.H., and K.L. Gould. 2003. The PCH family protein, Cdc15p, recruits two F-actin nucleation pathways to coordinate cytokinetic actin ring formation in Schizosaccharomyces pombe. Journal of Cell Biology. 162:851–862.

Chang, F., A. Woollard, and P. Nurse. 1996. Isolation and characterization of fission yeast mutants defective in the assembly and placement of the contractile actin ring. J Cell Sci. 109 (Pt 1):131–142.

Collins, A., A. Warrington, K.A. Taylor, and T. Svitkina. 2011. Structural organization of the actin cytoskeleton at sites of clathrin-mediated endocytosis. Curr Biol. 21:1167–1175.

Engqvist-Goldstein, A.E., and D.G. Drubin. 2003. Actin assembly and endocytosis: from yeast to mammals. Annu Rev Cell Dev Biol. 19:287–332.

Fernandez-Ballester, G., P. Beltrao, J.M. Gonzalez, Y.H. Song, M. Wilmanns, A. Valencia, and L. Serrano. 2009. Structure-based prediction of the Saccharomyces cerevisiae SH3-ligand interactions. J Mol Biol. 388:902–916.

Forsburg, S.L. 1993. Comparison of Schizosaccharomyces pombe expression systems. Nucleic Acids Res. 21:2955–2956.

Galletta, B.J., D.Y. Chuang, and J.A. Cooper. 2008. Distinct Roles for Arp2/3 Regulators in Actin Assembly and Endocytosis. PLoS Biology. 6.

Gibson, D.G., L. Young, R.Y. Chuang, J.C. Venter, C.A. Hutchison, 3rd, and H.O. Smith. 2009. Enzymatic assembly of DNA molecules up to several hundred kilobases. Nature Methods. 6:343–345.

Goode, B.L., J.A. Eskin, and B. Wendland. 2015. Actin and Endocytosis in Budding Yeast. Genetics. 199:315–358.

Higgs, H.N., and T.D. Pollard. 2001. Regulation of actin filament network formation through ARP2/3 complex: activation by a diverse array of proteins. Annu Rev Biochem. 70:649–676.

Huang, J., Y. Huang, H. Yu, D. Subramanian, A. Padmanabhan, R. Thadani, Y. Tao, X. Tang, R. Wedlich-Soldner, and M.K. Balasubramanian. 2012. Nonmedially assembled F-actin cables incorporate into the actomyosin ring in fission yeast. Journal of Cell Biology. 199:831–847.

Idrissi, F.Z., A. Blasco, A. Espinal, and M.I. Geli. 2012. Ultrastructural dynamics of proteins involved in endocytic budding. Proc Natl Acad Sci U S A. 109:E2587–2594.

Idrissi, F.Z., H. Grotsch, I.M. Fernandez-Golbano, C. Presciatto-Baschong, H. Riezman, and M.I. Geli. 2008. Distinct acto/myosin-I structures associate with endocytic profiles at the plasma membrane. Journal of Cell Biology. 180:1219–1232.

Kaksonen, M., and A. Roux. 2018. Mechanisms of clathrin-mediated endocytosis. Nat Rev Mol Cell Biol. 19:313–326.

Kaksonen, M., Y. Sun, and D.G. Drubin. 2003. A pathway for association of receptors, adaptors, and actin during endocytic internalization. Cell. 115:475–487.

Kaksonen, M., C.P. Toret, and D.G. Drubin. 2005. A modular design for the clathrin-and actin-mediated endocytosis machinery. Cell. 123:305–320.

Kaksonen, M., C.P. Toret, and D.G. Drubin. 2006. Harnessing actin dynamics for clathrin-mediated endocytosis. Nat Rev Mol Cell Biol. 7:404–414.

Keeney, J.B., and J.D. Boeke. 1994. Efficient targeted integration at leu1-32 and ura4-294 in Schizosaccharomyces pombe. Genetics. 136:849–856.

Kishimoto, T., Y. Sun, C. Buser, J. Liu, A. Michelot, and D.G. Drubin. 2011. Determinants of endocytic membrane geometry, stability, and scission. Proc Natl Acad Sci U S A. 108:E979–988.

Kovar, D.R., V. Sirotkin, and M. Lord. 2011. Three’s company: The fission yeast actin cytoskeleton. Trends in Cell Biology. 21:177–187.

Kukulski, W., M. Schorb, M. Kaksonen, and J.A. Briggs. 2012. Plasma membrane reshaping during endocytosis is revealed by time-resolved electron tomography. Cell. 150:508–520.

Laporte, D., V.C. Coffman, I.J. Lee, and J.Q. Wu. 2011. Assembly and architecture of precursor nodes during fission yeast cytokinesis. Journal of Cell Biology. 192:1005–1021.

Le Goff, X., F. Motegi, E. Salimova, I. Mabuchi, and V. Simanis. 2000. The S. pombe rlc1 gene encodes a putative myosin regulatory light chain that binds the type II myosins myo3p and myo2p. Journal of Cell Science. 113 Pt 23:4157–4163.

Lee, W.L., M. Bezanilla, and T.D. Pollard. 2000. Fission yeast myosin-I, Myo1p, stimulates actin assembly by Arp2/3 complex and shares functions with WASp. Journal of Cell Biology. 151:789–800.

Maundrell, K. 1990. nmt1 of fission yeast. A highly transcribed gene completely repressed by thiamine. Journal of Biological Chemistry. 265:10857–10864.

McMahon, H.T., and E. Boucrot. 2011. Molecular mechanism and physiological functions of clathrin-mediated endocytosis. Nat Rev Mol Cell Biol. 12:517–533.

Michelot, A., M. Costanzo, A. Sarkeshik, C. Boone, J.R.Y. III, and D.G. Drubin. 2010. Reconstitution and Protein Composition Analysis of Endocytic Actin Patches. Current Biology. 20:1890–1899.

Mochida, J., T. Yamamoto, K. Fujimura-Kamada, and K. Tanaka. 2002. The novel adaptor protein, Mti1p, and Vrp1p, a homolog of Wiskott-Aldrich syndrome protein-interacting protein (WIP), may antagonistically regulate type I myosins in Saccharomyces cerevisiae. Genetics. 160:923–934.

Mund, M., J.A.v.d. Beek, J. Deschamps, S. Dmitrieff, J.L. Monster, A. Picco, F. Nédélec, M. Kaksonen, and J. Ries. Nov. 15, 2017. Systematic analysis of the molecular architecture of endocytosis reveals a nanoscale actin nucleation template that drives efficient vesicle formation. bioRxiv.

Naqvi, S.N., R. Zahn, D.A. Mitchell, B.J. Stevenson, and A.L. Munn. 1998. The WASp homologue Las17p functions with the WIP homologue End5p/verprolin and is essential for endocytosis in yeast. Curr Biol. 8:959–962.

Pelham, R.J., Jr., and F. Chang. 2001. Role of actin polymerization and actin cables in actin-patch movement in Schizosaccharomyces pombe. Nature Cell Biology. 3:235–244.

Picco, A., W. Kukulski, H.E. Manenschijn, T. Specht, J.A.G. Briggs, and M. Kaksonen. 2018. The contributions of the actin machinery to endocytic membrane bending and vesicle formation. Mol Biol Cell. 29:1346–1358.

Picco, A., M. Mund, J. Ries, F. Nedelec, and M. Kaksonen. 2015. Visualizing the functional architecture of the endocytic machinery. Elife. 4.

Pollard, L.W., M. Onishi, J.R. Pringle, and M. Lord. 2012. Fission yeast Cyk3p is a transglutaminase-like protein that participates in cytokinesis and cell morphogenesis. Mol Biol Cell. 23:2433–2444.

Ramesh, N., I.M. Anton, J.H. Hartwig, and R.S. Geha. 1997. WIP, a protein associated with wiskott-aldrich syndrome protein, induces actin polymerization and redistribution in lymphoid cells. Proc Natl Acad Sci U S A. 94:14671–14676.

Roberts-Galbraith, R.H., J.S. Chen, J. Wang, and K.L. Gould. 2009. The SH3 domains of two PCH family members cooperate in assembly of the Schizosaccharomyces pombe contractile ring. Journal of Cell Biology. 184:113–127.

Roberts-Galbraith, R.H., M.D. Ohi, B.A. Ballif, J.S. Chen, I. McLeod, W.H. McDonald, S.P. Gygi, J.R. Yates, 3rd, and K.L. Gould. 2010. Dephosphorylation of F-BAR protein Cdc15 modulates its conformation and stimulates its scaffolding activity at the cell division site. Molecular Cell. 39:86–99.

Rodal, A.A., A.L. Manning, B.L. Goode, and D.G. Drubin. 2003. Negative regulation of yeast WASp by two SH3 domain-containing proteins. Current Biology. 13:1000–1008.

Rodal, A.A., O. Sokolova, D.B. Robins, K.M. Daugherty, S. Hippenmeyer, H. Riezman, N. Grigorieff, and B.L. Goode. 2005. Conformational changes in the Arp2/3 complex leading to actin nucleation. Nature Structural & Molecular Biology. 12:26–31.

Rosenberg, J.A., G.C. Tomlin, W.H. McDonald, B.E. Snydsman, E.G. Muller, J.R. Yates, 3rd, and K.L. Gould. 2006. Ppc89 links multiple proteins, including the septation initiation network, to the core of the fission yeast spindle-pole body. Molecular Biology of the Cell. 17:3793–3805.

Sato, M., S. Dhut, and T. Toda. 2005. New drug-resistant cassettes for gene disruption and epitope tagging in Schizosaccharomyces pombe. Yeast. 22:583–591.

Shaner, N.C., G.G. Lambert, A. Chammas, Y. Ni, P.J. Cranfill, M.A. Baird, B.R. Sell, J.R. Allen, R.N. Day, M. Israelsson, M.W. Davidson, and J. Wang. 2013. A bright monomeric green fluorescent protein derived from Branchiostoma lanceolatum. Nature Methods. 10:407–409.

Siam, R., W.P. Dolan, and S.L. Forsburg. 2004. Choosing and using Schizosaccharomyces pombe plasmids. Methods. 33:189–198.

Sirotkin, V., C.C. Beltzner, J.B. Marchand, and T.D. Pollard. 2005. Interactions of WASp, myosin-I, and verprolin with Arp2/3 complex during actin patch assembly in fission yeast. Journal of Cell Biology. 170:637–648.

Sirotkin, V., J. Berro, K. Macmillan, L. Zhao, and T.D. Pollard. 2010. Quantitative analysis of the mechanism of endocytic actin patch assembly and disassembly in fission yeast. Molecular Biology of the Cell. 21:2894–2904.

Skau, C.T., and D.R. Kovar. 2010. Fimbrin and tropomyosin competition regulates endocytosis and cytokinesis kinetics in fission yeast. Curr Biol. 20:1415–1422.

Skruzny, M., T. Brach, R. Ciuffa, S. Rybina, M. Wachsmuth, and M. Kaksonen. 2012. Molecular basis for coupling the plasma membrane to the actin cytoskeleton during clathrin-mediated endocytosis. Proc Natl Acad Sci U S A. 109:E2533–2542.

Snaith, H.A., I. Samejima, and K.E. Sawin. 2005. Multistep and multimode cortical anchoring of tea1p at cell tips in fission yeast. EMBO J. 24:3690–3699.

Sohrmann, M., C. Fankhauser, C. Brodbeck, and V. Simanis. 1996. The dmf1/mid1 gene is essential for correct positioning of the division septum in fission yeast. Genes Dev. 10:2707–2719.

Soulard, A., T. Lechler, V. Spiridonov, A. Shevchenko, A. Shevchenko, R. Li, and B. Winsor. 2002. Saccharomyces cerevisiae Bzz1p is implicated with type I myosins in actin patch polarization and is able to recruit actin-polymerizing machinery in vitro. Molecular Biology of the Cell. 22:7889–7906.

Stark, B.C., M.L. James, L.W. Pollard, V. Sirotkin, and M. Lord. 2013. UCS protein Rng3p is essential for myosin-II motor activity during cytokinesis in fission yeast. PLoS One. 8:e79593.

Sun, Y., A.C. Martin, and D.G. Drubin. 2006. Endocytic internalization in budding yeast requires coordinated actin nucleation and myosin motor activity. Developmental Cell. 11:33–46.

Tong, A.H.Y., B. Drees, G. Nardelli, G.D. Bader, B. Brannetti, L. Castagnoli, M. Evangelista, S. Ferracuti, B. Nelson, S. Paoluzi, M. Quondam, A. Zucconi, C.W.V. Hogue, S. Fields, C. Boone, and G. Cesareni. 2002. A Combined Experimental and Computational Strategy to Define Protein Interaction Networks for Peptide Recognition Modules. Science. 295.

Tonikian, R., X. Xin, C.P. Toret, D. Gfeller, C. Landgraf, S. Panni, S. Paoluzi, L. Castagnoli, B. Currell, S. Seshagiri, H. Yu, B. Winsor, M. Vidal, M.B. Gerstein, G.D. Bader, R. Volkmer, G. Cesareni, D.G. Drubin, P.M. Kim, S.S. Sidhu, and C. Boone. 2009. Bayesian modeling of the yeast SH3 domain interactome predicts spatiotemporal dynamics of endocytosis proteins. PLoS Biol. 7:e1000218.

Wach, A., A. Brachat, R. Pohlmann, and P. Philippsen. 1994. New heterologous modules for classical or PCR-based gene disruptions in Saccharomyces cerevisiae. Yeast. 10:1793–1808.

Wagner, A.R., Q. Luan, S.-L. Liu, and B.J. Nolen. 2013. Dip1 Defines a Class of Arp2/3 Complex Activators that Function without Preformed Actin Filaments. Current Biology. 23:1990–1998.

Weinberg, J., and D.G. Drubin. 2012. Clathrin-mediated endocytosis in budding yeast. Trends in Cell Biology. 22:1–13.

Willet, A.H., N.A. McDonald, K.A. Bohnert, M.A. Baird, J.R. Allen, M.W. Davidson, and K.L. Gould. 2015. The F-BAR Cdc15 promotes contractile ring formation through the direct recruitment of the formin Cdc12. Journal of Cell Biology. 208:391–399.

Wu, J.Q., J.R. Kuhn, D.R. Kovar, and T.D. Pollard. 2003. Spatial and temporal pathway for assembly and constriction of the contractile ring in fission yeast cytokinesis. Developmental Cell. 5:723–734.

Wu, J.Q., V. Sirotkin, D.R. Kovar, M. Lord, C.C. Beltzner, J.R. Kuhn, and T.D. Pollard. 2006. Assembly of the cytokinetic contractile ring from a broad band of nodes in fission yeast. Journal of Cell Biology. 174:391–402.

Yakura, M., F. Ozoe, H. Ishida, T. Nakagawa, K. Tanaka, H. Matsuda, and M. Kawamukai. 2006. zds1, a novel gene encoding an ortholog of Zds1 and Zds2, controls sexual differentiation, cell wall integrity and cell morphology in fission yeast. Genetics. 172:811–825.

Young, M.E., J.A. Cooper, and P.C. Bridgman. 2004. Yeast actin patches are networks of branched actin filaments. Journal of Cell Biology. 166:629.

